# A Glutamine antagonist-modulated tumor microenvironment unleashes enzalutamide’s immunotherapeutic effects

**DOI:** 10.64898/2026.06.30.735585

**Authors:** Jianpeng Yu, Xue Jiang, Haipei Yao, Zichun Xing, Fan Zhang, Chen Jin, Moyasar A. Alhamo, Hong Zhang, Bangchen Wang, Michelle L. Bowie, Olivia Meng, Daniel J. George, Robert Wild, Xia Gao, Yi Zhang, David M Ashley, Christopher J Pirozzi, Herman F. Staats, Yiping He, Jiaoti Huang

**Author notes:** Corresponding: Jiaoti Huang, MD, PhD, Yiping He, PhD.

## Abstract

Androgen receptor (AR) antagonists, such as enzalutamide, suppress prostate cancer (PCa) cells to achieve temporary therapeutic effects. In addition to tumor cell-autonomous suppressive function, AR antagonists can also potentially exert anti-tumor effects via mitigating cytotoxic T cells’ exhaustion. However, strategies for effectively harnessing enzalutamide’s immunotherapeutic effects remain elusive. In studying a recently described glutamine antagonist prodrug (DRP-104) in PCa models, we found that despite the initial response, tumors ultimately became resistant. Intriguingly, compared to the untreated (DRP-104 treatment-naïve) tumors, the resistant tumors became highly susceptible to enzalutamide in vivo. Additionally, treating tumors with DRP-104 and enzalutamide simultaneously also yielded superior therapeutic efficacy. We demonstrated that DRP-104 therapy promoted the infiltration of CD8^+^ T cells as well as regulatory T cells (Treg) in responsive tumors, and the tumor-infiltrating Treg were mostly depleted upon enzalutamide treatments. Mechanistically, we showed that Treg differentiation from mouse CD4^+^ T cells was attenuated by enzalutamide. We further demonstrated that Treg induction was accompanied by the interaction between AR and aryl hydrocarbon receptor (AhR), the nuclear receptor indispensable for Treg differentiation, in the nuclei of CD4^+^ T cells, and this interaction was diminished by enzalutamide. In further support of AR signaling in Treg biogenesis, analysis of available gene expression datasets found that *AR* expression was elevated in Treg when compared to CD4^+^ T cells in human peripheral blood mononuclear cells (PBMCs). In addition, it was positively correlated with Treg module scores in several human cancer types. Finally, using an anti-GPC3 (Glypican 3) vaccination model, we demonstrated that CD4^+^ T cells subjected to Treg induction in the presence of enzalutamide were less effective in protecting GPC3-expressing tumor cells from CD8^+^ T cell’s cytotoxic killing. Collectively, these results suggest that AR promotes Treg’ s differentiation and/or immunosuppressive functions, and nominate enzalutamide as a Treg-mitigating agent for potentiating immunotherapies. Our results also demonstrate that an otherwise unintended, Treg-promoting property of DRP-104 can be leveraged to unleash the immune-regulatory function of enzalutamide for the treatment of PCa.

## Introduction

AR is the central driver of and the most extensively studied therapeutic target in PCa. As a majority of PCas are histologically classified as AR-dependent adenocarcinomas, AR-targeting therapies (including androgen deprivation therapy and AR antagonists such as enzalutamide) are the mainstays for clinical management of PCa. Additionally, research in other cancer types has revealed widespread dysregulation of the AR signaling, and points to context-dependent effects of AR on cancer pathogenesis and progression (1).

Despite the tumor cell-autonomous effects of the AR signaling, recent research has uncovered profound effects of AR in modulating the functions of tumor microenvironment, particularly T lymphocytes. An initial study focusing on PCa reveals that AR signaling contributes to T cell exhaustion in the tumors, leading to the finding that AR inhibition augments CD8^+^ T cell’s anti-tumor activity and can sensitize PCa to immune checkpoint blockade(2). Subsequently, this effect of AR signaling to T lymphocytes’ dysfunctions, particularly their exhaustion in cancers, has been observed in other cancer types, leading to the notion that the levels of androgens can partially explain sex-dependent differences in autoimmunity, cancers’ immune landscapes, and their responses to immunotherapies (3–6).

These independent lines of evidence support that AR antagonists can be leveraged to mitigate CD8^+^ T cell exhaustion to achieve more effective cancer immunotherapies(2–4). However, enzalutamide, the AR-targeting drug designed to directly suppress AR-dependent prostate cancer cells, has been associated with enhanced immunosuppression in recurrent PCa(7, 8). In addition, combining enzalutamide with checkpoint blockade has failed to yield clinical benefits in PCa patients(9). Thus, the benefits of this standard-of-care PCa drug, or other AR antagonists commonly used for treating PCa, in the context of PCa immunotherapy remain uncertain.

We previously demonstrated that dysregulated glutamine metabolism is a major driver of PCa progression, and targeting glutamine utilization can be an effective therapeutic strategy(10, 11). Subsequently, we reported potent anti-tumor effects of DRP-104, a glutamine antagonist prodrug, in PCa, via both tumor cell-autonomous suppression and stimulating the host’s immune system(12, 13). Specifically, we showed that in preclinical PCa models, blocking glutamine utilization effectively suppresses PCa tumors while inducing infiltration of CD8^+^ T lymphocytes into the tumor tissue(13).

Here, using PCa models, we show that despite promising anti-tumor efficacy of the glutamine blockade strategy, cancer cells ultimately develop resistance to the treatment. These resistant cells display a similar response to AR inhibition (i.e., hormone depletion or enzalutamide) when compared to their untreated (DRP-104-naïve) counterparts in in vitro assays. Surprisingly, we found that the recurrent (resistant) tumors became more sensitive to enzalutamide compared to their treatment-naïve counterparts. This enhanced efficacy was also observed when naïve tumors were treated with the dual agents simultaneously. Mechanistically, we showed that upon monotherapy with DRP-104, the responsive tumors were infiltrated with CD8^+^ T cells as well as Treg. We discovered that while enzalutamide as a monotherapy failed to induce T cell infiltration in naïve tumors, it promoted the presence of CD8^+^ T cells when combined with DRP-104, and rendered DRP-104-resistant tumors free of Treg. Our results suggest that enzalutamide can enable anti-tumor immune response and curtail the differentiation and/or function of Treg, thus providing a mechanism underlying its potency against the DRP-104-treated recurrent tumors. Mechanistic and gene expression analyses implicate AR as a critical factor in controlling Treg differentiation and/or function and in shaping Treg homeostasis in PCa.

## Methods and Materials

### Cell lines, transfection and stable cell line establishment

Prostate cancer cell lines (LNCaP, C4-2, PC3 and TrampC2) were purchased from the American Type Culture Collection (ATCC) via Duke Cell Culture Facility (CCF). The cell lines’ short tandem repeat profiles were confirmed by Duke CCF, and the cell lines were tested periodically to ensure that they were *Mycoplasma*-free. All human prostate cancer cell lines were cultured as previously described (12). PPR, an NEPC mouse cell line, was a gift from Dr. Ming Chen (14). Briefly, LNCaP, C4-2 and PC3 cells were maintained in RPMI 1640 medium with L-glutamine (Gibco, 11875093, ThermoFisher, Waltham, MA, USA), supplemented with 10% fetal bovine serum and 1% penicillin-streptomycin. Charcoal dextran stripped fetal bovine serum (CSS) (Denville) treatment of LNCaP was carried out by using RPMI 1640 (Gibco, 11835, phenol red free), supplemented with 10% CSS and 1% penicillin-streptomycin as previously described (11). TrampC2 cells were maintained in DMEM with L-glutamine (Gibco, 11965092, ThermoFisher), supplemented with 10% fetal bovine serum, 0.005 mg/ml human insulin, and 10 nM dehydroisoandrosterone and 1% penicillin-streptomycin. PPR cells were maintained in DMEM, supplemented with 10% fetal bovine serum, 0.005 mg/ml bovine insulin, 25 µg/mL bovine pituitary extract, 6 ng/mL recombinant human EGF, 1 nM dihydrotestosterone, and 1% penicillin-streptomycin. DRP-104-resistant cells were generated in our previous study (13). TrampC2 cells were seeded in 6-cm dishes and transfected with the pCMV6-GPC3 plasmid (MC219262, OriGene Technologies, USA) using Lipofectamine 3000 (Invitrogen, L3000015, Thermo Fisher Scientific) according to the manufacturer’s instructions at a DNA-to-reagent ratio of 1:1.5. Stable transfectants were selected with 500 μg/mL G418 for 7 days, followed by maintenance in 300 μg/mL G418 for an additional 7 days.

### Chemicals and antibodies

DRP-104 was previously described (13). Enzalutamide was obtained from MedChemExpress (cat# HY-70002). Antibodies used for immunohistochemistry staining included the following: CD8α (1:100 dilution, A02236-1, Boster Bio), FOXP3 (1:100 dilution, RP1032, Boster Bio; 1:100 dilution, 65089-1-Ig, Proteintech), CgA (1:500 dilution, 10529-1-AP, Proteintech), PEG10(1:500 dilution, 14412-1-AP, Proteintech). Antibodies used for immunofluorescence (IF) staining and PLA assay included the following: CD4 (20 µg/ml, 50-9766-82, Invitrogen), FOXP3 (1:100 dilution, RP1032, Boster Bio; 1:100 dilution, 65089-1-Ig, Proteintech), GPC3 (2 µg/ml, 222514-rAb, Addgene), AR (1:200 dilution, 22089-1-AP, Proteintech), AHR (1:200 dilution, 67785-1-Ig, Proteintech). Antibodies used for flow cytometry included the following: CD4 (20 µg/ml, 50-9766-82, Invitrogen), FOXP3 (2 µg/ml, 560408, BD Pharmingen).

### In vitro cell growth assays

In vitro cell growth assays were performed following a previously described procedure (13). Briefly, adherent cells were plated onto a 96-well plate at ∼5% confluency with at least 3 biological replicates. Cell growth was monitored and quantified by confluency using the IncuCyte S3 Live-Cell Analysis system (Sartorius, Göttingen, Germany, RRID:SCR_023147) with nine images per well taken with a 20x objective. The change in confluency was normalized to the confluency at Day 0. All cell growth assays were repeated in at least two independent experiments.

### Metabolite profiling

C4-2 and PC3 metabolite profiling data were retrieved from our previous study (13). Briefly, cells were plated into six-well plates at 35% confluency with three technical replicates and treated with DMSO (vehicle), 5 μmol/L DRP-104 for 48 hours. The confluency of each of the wells was measured using the IncuCyte S3 Live-Cell Analysis system (Sartorius) before metabolite isolation. Cellular metabolites were extracted, as described in our previous study (13). Cells were collected in 80% methanol to extract the metabolites. After centrifugation, the supernatant was dried with a speed vacuum, and dry pellets were reconstituted into 30 μL of sample solvent (water:methanol:acetonitrile, 2:1:1), followed by centrifugation at 20,000 × g for 3 minutes. The supernatant was transferred to liquid chromatography vials, and 3 μL was submitted to be analyzed with high-performance LC/MS.

### In vivo tumor models and treatments

The animal study was approved by the Institutional Animal Care and Use Committee of Duke University. All mice were purchased from The Jackson Laboratory. For TrampC2- and PPR-derived tumor models, immune-competent (wild-type) 6 to 8-week-old C57BL/6J (JAX strain #000664) mice were used. The subcutaneous inoculation was performed by injecting 0.5 to 1 million cells in PBS and Matrigel (Corning) mixture (4:1) under the right flank skin using a 27-gauge × 0.5-inch needle. This injection was performed under brief isoflurane anesthesia on a shaved ∼1-cm^2^ region of the flank. When the tumors reached approximately 50 to 500 mm^3^, mice were randomized into 5 to 8 mice per group and treated via subcutaneous and intraperitoneal injections per the following treatments: (i) Tween 80: ethanol: saline (5:5:90) vehicle (subcutaneous injection) and corn oil vehicle (intraperitoneal injection); (ii) Tween 80:ethanol:saline (5:5:90) vehicle (subcutaneous injection) and 10 mg/kg Enzalutamide (intraperitoneal injection); (iii) 2 mg/kg DRP-104 (subcutaneous injection) and corn oil vehicle (intraperitoneal injection); or (iv) 2 mg/kg DRP-104 (subcutaneous injection) and 10 mg/kg Enzalutamide (intraperitoneal injection). Xenograft sizes were measured every 2 to 4 days using calipers by the same scientist throughout the experiment and calculated using the formula tumor volume = length × (width)^2^ × 0.5. Mice were sacrificed when the tumor size met humane end-stage criteria. Syngeneic tumors were dissected and imaged.

### Hematoxylin and eosin staining and immunohistochemistry staining

Hematoxylin and eosin (H&E) and immunohistochemistry staining were performed following standard procedures, as we previously described (13, 15). Formalin-fixed, paraffin-embedded sections at 4–5 mm were deparaffinized and rehydrated. Heat-mediated antigen retrieval was performed in a boiled water bath for 40 minutes in citrate buffer (pH 6.0). Antibodies were incubated at room temperature for 1 hour. Then the slides were incubated for 45 minutes at room temperature with horseradish peroxidase–conjugated secondary antibodies (Dako EnVision + Kit) and visualized with 3,3’-diaminobenzidine (DAB).

### Reverse Transcription and Real-Time Quantitative PCR (RT-qPCR)

Total RNA was extracted from the cell pellet using an RNA extraction kit (cat. #R1055, Zymo Research, California, USA) following the manufacturer’s instructions. The RNA concentration and absorbance were measured using NanoDrop 2000 (Thermo Fisher Scientific, RRID: SCR_018042). RNA samples with a concentration exceeding 15 ng/μL and an A260/A280 ratio above 1.9 were used for reverse transcription to synthesize cDNA (cat. #11155ES60, Yeasen Biotechnology, Maryland, USA). SYBR Green qPCR Master Mix (cat. #11184ES08, Yeasen Biotechnology) and corresponding primers were mixed and added to each reaction well, with each target gene analyzed in Triplicate. RT-qPCR was performed on a QuantStudio 3 instrument (Thermo Fisher Scientific, RRID: SCR_018712). The housekeeping gene Actb was used for normalization, and the relative expression of target genes was calculated using the 2^−ΔΔCT^ method. Primers used for PCR:

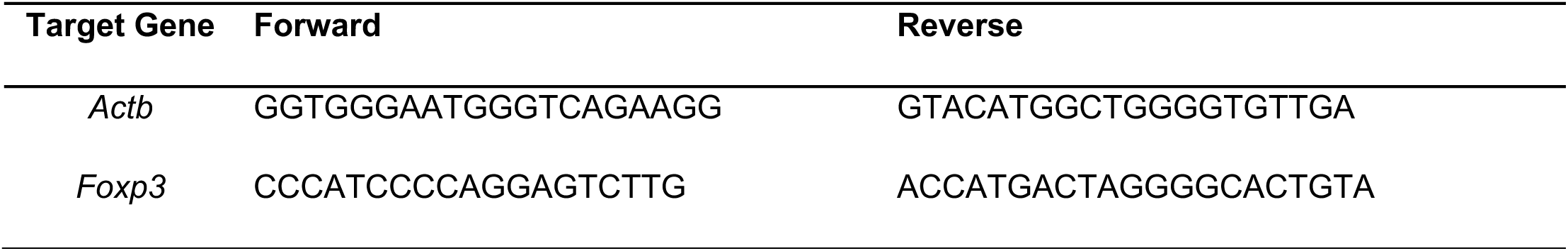

### Immunofluorescent staining and proximity ligation assay

For CD4 and FOXP3 immunofluorescent staining, formalin-fixed, paraffin-embedded sections at 4 mm were deparaffinized and rehydrated. Heat-mediated antigen retrieval was performed in a boiled water bath for 40 minutes in citrate buffer (pH 6.0). Anti-CD4 with fluorescence label and anti-FOXP3 were incubated at room temperature for 1 hour. Then the slides were incubated for 45 minutes at room temperature with the secondary antibody for anti-FOXP3 (anti-rabbit Alexa 488, 1:500 dilution, Thermo Fisher Scientific, Waltham, MA, USA). Subsequently, slides were washed and stained with 1 μg/mL DAPI (MilliporeSigma, #D9542, Darmstadt, Germany) before the slides were mounted with ProLong Gold Antifade Mountant (Thermo Fisher Scientific, #P36394, Waltham, MA, USA). The slides were imaged using the Zeiss 880 Airyscan Fast Inverted Confocal Microscope and analyzed using Zeiss ZEN 3.9 (blue edition) software (Carl Zeiss Microscopy GmbH).

For GPC3 immunofluorescent staining, cells were washed with PBS, fixed in 4% formaldehyde (Santa Cruz Biotechnology) at room temperature for 15 minutes, and permeabilized with 0.2% Triton X-100 (Bio-Rad) for 15 minutes. Slides with fixed and permeabilized cells were blocked with blocking buffer (1% BSA, 0.2% Triton X-100, 10% goat serum, 0.2 M glycine in PBS) at room temperature for 1 hour and then incubated with primary antibody diluted in antibody dilution buffer (1% BSA, 0.2% Triton X-100 in PBS) at room temperature for 1 hour. Then, the slides were washed three times with PBS and incubated with the secondary antibody (anti-rabbit Alexa 488, 1:500 dilution, Thermo Fisher Scientific) at room temperature for 45 minutes. Afterwards, the cells were washed, stained with 1 µg/mL DAPI (MilliporeSigma), and mounted on slides using ProLong Gold Antifade Mountant (ThermoFisher). The slides were imaged with Zeiss 880 Airyscan Fast Inverted Confocal Microscope and analyzed using the Zeiss Zen 3.9 (blue edition) software (Carl Zeiss Microscopy GmbH).

The proximity ligation assay (PLA) was conducted following the manufacturer’s protocol (MilliporeSigma, DUO92102-1KT). In brief, cells were washed with PBS and fixed with 4% paraformaldehyde (PFA) in PBS at room temperature for 15 minutes. Subsequently, the fixed cells were rinsed with PBS. Permeabilization was carried out with 0.2% Triton X-100 in PBS at room temperature for 10 minutes, followed by washing with PBS. The cells were then blocked with PLA blocking solution at room temperature for 1 hour. Primary antibodies, diluted in PLA diluent, were incubated with the cells at room temperature for 1 hour. After removing the primary antibody solution, cells were washed twice with PLA wash buffer A at room temperature and incubated with the PLUS and MINUS PLA probes. Next, PLA ligation solution was prepared and incubated with cells at 37 °C in a humidified incubator for 30 minutes, followed by two washes with PLA wash buffer A at room temperature. Then, PLA amplification solution was prepared and incubated with cells at 37 °C for 100 minutes. Following the incubation, cells were washed twice with PLA wash buffer B. The cell nuclei was stained with DAPI dye. Finally, the PLA sample slides were assembled and ready for imaging.

### RNA sequencing and analysis

Tumor cells or tissues were harvested, washed twice with ice-cold PBS, and total RNA was extracted using the RNeasy Mini Kit (Qiagen, 74104, Maryland, USA) following the manufacturer’s protocol. RNA samples were subjected to DNase treatment and quality assessment prior to mRNA capture and standard mRNA sequencing (paired-end, 150 bp sequencing) on an Illumina NovaSeq 6000 or NovaSeq X Plus platform (Genewiz, Azenta Life Sciences City, NJ).

For transcriptomic profiling for mouse T cells, total RNA was used as the starting material. Watchmaker RNA Library Prep Kit with Polaris™ Depletion (7BK0002, Watchmaker Genomics, USA) was used to prepare libraries. Briefly, rRNA was removed from total RNA by using mouse rRNA-specific probes. Fragmentation was carried out using divalent cations under elevated temperature in First Strand Synthesis Reaction Buffer. First, strand cDNA was synthesized using random hexamer primer and reverse transcriptase. Then the second strand of cDNA was synthesized by adding buffer and dNTPs (dTTP in dNTP is replaced by dUTP). The synthesized double-stranded cDNA was used for end repair, A-tailing, adaptor ligation, fragment selection and PCR amplification. PCR products were purified to obtain a strand specific library. After the library construction was completed, preliminary quantification was carried out using Qubit. Subsequently, the inserted fragments of the library were detected. Once the inserted fragments met desired expectations, the effective concentration of the library was accurately quantified using qRT-PCR. The libraries were pooled and sequenced on Illumina platforms, according to effective library concentration and data amount required.

Raw sequencing reads underwent quality control using FastQC (Galaxy Version 0.74+galaxy1) to assess base quality scores and detect potential sequencing artifacts. Sequencing reads were aligned (0-500 fragment length for valid paired-end alignments) to the human reference genome (GRCh37/hg19) or Mus musculus genome (GRCm38/mm10) using Bowtie2 (Galaxy Version 2.5.3+galaxy1) with default parameters. Differential gene expression analysis was conducted using DESeq2 with outliers filtering (Galaxy Version 2.11.40.8+galaxy0). Principal component analysis (PCA) was performed with a convex type of ellipses by using PCA plot w ggplot2 (Galaxy Version 3.4.0+galaxy1) (doi:10.1093/nar/gkae410). Gene Set Enrichment Analysis (GSEA) was conducted with GSEA (4.1.0) (16, 17). GO Enrichment analysis was performed with ShinyGO 0.85.1(18). The mouse transcriptomics deconvolution was performed with conversion of murine genes to human genes through orthologous gene mapping by using mouse2human online tool (19–22) and quantified using immunedeconv with quanTIseq implemented (23). The enrichment analysis for immune cell types across the T cell sequencing samples were performed with Cross-tissue Immune cell type or state Enrichment analysis of gene lists for Cancer (CIEC) online tool (24).

### Treg Differentiation from CD4^+^ T cells

Splenocytes from C57BL/6J male mice (6–12-week-old) were harvested from the host after euthanasia, and the tissue was broken into single cells by pulverizing it into a 40 μm filter over a 50 mL conical tube. DPBS- and 2% FBS was then perfused through the filter to acquire remaining cell material before centrifuging the cell suspension at 600 g for 5 minutes at room temperature. Splenocytes were then treated with ACK Lysing Buffer (Cat.#: A1049201, ThermoFisher) for five minutes before resuspension for downstream culture. The cells were then counted and negatively selected for CD4^+^ T cells using the CD4^+^ T cell isolation kit by Stem Cell Technologies (Cat.#: 19852) and the manufacturer’s instructions. CD4^+^ T cells were then seeded into cell culture plates that had been previously coated in αCD3 antibody (Cat.#: 16-0031-85, ThermoFisher) overnight at 4°C at a concentration of 1 μg/mL. RPMI 1640 Complete media (10% FBS, 1% Anti-Anti, 10 mM HEPES, 1mM sodium pyruvate, 100 μM non-essential amino acids, and 50 μM beta-mercaptoethanol) was supplemented with T regulatory cell supplement (10957, STEMCELL Technologies) at a 1:100 ratio and 0.5 μg/mL of αCD28 (553295, BD Pharmingen). Alternatively, for Treg differentiation, above T cell complete media were supplemented with PC3 tumor cell-conditioned media (8 ml media in 10-cm plate with cells 70-80% confluency; 3-day cultured condition media without or with DRP-104 at 4 μM), IL2 (2 ng/ml, 212-12-5UG, Gibco), TGF-β (50 ng/ml, 7666-MB-005/CF, R&D Systems, Minnesota, USA), and *all−trans*−Retinoic acid (ATRA) (1 μM, R2625, MilliporeSigma). Cells were kept under sterile culture conditions at 37°C for 6 days before harvest.

### Analysis of Treg via FACS (Fluorescence-Activated Cell Sorting) Staining

For analysis of Treg, mouse CD4^+^ T cells were stained using antibodies directed at Foxp3. Single cell suspensions were pre-incubated (15 minutes on ice) with anti-CD16/32 Fc block and washed 3 times in FACS staining buffer (00-4222-26, eBioscience) before staining with anti-Foxp3 antibody conjugated with eFluor® 660 and incubating for 60 minutes on ice in the dark. Then the cells were washed, fixed, permeabilized, and stained following the eBioscience™ Foxp3 / Transcription Factor Staining Buffer Set protocol (00-5523-00, eBioscience) with 1-hour FOXP3 staining at room temperature. Standard flow cytometric techniques were used to acquire data on a 3-laser FACSCanto II (BD Biosciences). Analysis was performed using FlowJo (10.8.1, Becton Dickinson & Company) software.

### Vaccination using GPC3-derived Peptides

Nine mouse GPC3 protein-derived peptides predicted to be T cell epitopes in C57BL/6J mice by the Immune Epitope Database (IEDB)were selected for screening to identify peptides with immunogenicity(25). For immunization, mice under isoflurane anesthesia were intramuscularly injected with 100 μg of the indicated mixed peptides and adjuvants (25 μL of Addavax (vac-adx-10, InvivoGen, CA, USA) and 20 μg of CpG 1826 (HY-146245, MedChemExpress)) in sterile PBS in a total volume of 50 μl. Two or three vaccinations were performed before splenocytes were harvested for subsequent assays.

### IFN-γ enzyme-linked immunospot assay (ELISpot)

ELISpot was performed using splenocytes isolated from mice after the indicated vaccinations. The mouse IFN-γ ELISpot assay was performed using the R&D Systems™ Mouse IFN-γ ELISpot Kit (XEL485, R&D Systems) according to the manufacturer’s protocol. Briefly, Splenocytes (2.5 × 10^5^ or 5 × 10^5^ cells/well) were spread in the 96-well polyvinylidene fluoride–backed microplate coated with a monoclonal antibody specific for mouse IFN -γ. For the screening assay, the Splenocytes (2.5 × 10^4^ or 5 × 10^4^ cells/well) were spread in the 96-well microplate. Splenocytes were stimulated with 5 μg/ml GPC3 peptides. Tests were conducted with a negative control containing only medium and a positive control (Concanavalin A Solution, 00-4978-93, eBioscience). Cells were incubated in a 37 °C incubator for 24 hours. Afterward, cells were removed, and the plates were washed with the wash buffer supplied by the kit, and incubated with the diluted Detection Antibody mixture overnight at 4 °C. After washing, the plates were incubated with streptavidin conjugated to alkaline phosphatase for 2 hours at room temperature. Incubated plates were developed using BCIP/NBT Substrate for 1 hour at room temperature. Spots were counted using an ImmunoSpot reader and performed by the Regional Biocontainment Laboratory (RBL) at Duke University.

### Cytotoxic T lymphocyte (CTL) assay

Re-stimulation of effector cells. C57BL/6J splenocytes were resuspended and cultured in RPMI 1640 Complete media (10% FBS, 1% Anti-Anti, 10 mM HEPES, 1mM sodium pyruvate, 100 μM non-essential amino acids, and 50 μM beta-mercaptoethanol) with the addition of IL2 (20 ng/ml, 212-12-5UG, Gibco) and CTL epitope peptides at 1 μg/ml (w/o) for 6 days. On day 3, the equal volume fresh media were added to each well. CD8^+^ cells were isolated with Mouse CD8+ T Cell Isolation Kit (19853, Stemcell Technologies) after restimulation for 6 days and cocultured with tumor cells, w/o induced Tregs for 2 days. A standard CellTiter-Glo® Luminescent Cell Viability (CTG) Assay (G7572, Promega, WI, USA) with minor modification was used to monitor CTL activity. Results are presented as peptide-specific lysis at the Effector-to-Target (E: T) ratio of 20:1.

### Analysis of public datasets

The Database of Immune Cell Expression (DICE) was used to analyze the expression of *AR* in human PBMCs(26). Three publicly available scRNA-seq datasets were used to assess the correlations between the cell-state/subtype (exhausted CD8^+^ T cell and effector Treg) signatures and AR signal activity in tumors, including GSM4972161 (glioma)(27), GSE139324 (head and neck cancer)(28), and GSE150430 (nasopharyngeal carcinoma)(29). AR activity was calculated by the module score of the human gene set Hallmark_Androgen_Response(30). For the glioma dataset, CD4^+^ and CD8^+^ T cells were annotated by the original study (GSE163108)(27). For the head and neck cancer and nasopharyngeal carcinoma datasets, CD4^+^ and CD8^+^ T cells were defined by positive *CD4* expression or positive *CD8A* expression, respectively. The effector Treg signature was defined by the expression of *FOXP3*, *CTLA-4*, *CCR8*, and *TNFRSF9* (31, 32). Exhausted CD8^+^ T cells were defined by the previously described 82-gene panel (33). All module scores were calculated by using sc.tl.score_genes function in scanpy (v.1.11.5) for every single cell, and were averaged across all cells within each patient for sample-level estimations(34). Spearman correlation coefficients between the sample-level cell-state/subtype (exhausted CD8^+^ T cell signature and effector Treg) signature scores and AR activity scores were calculated using scipy (v. 1.17.1)(35). For estimating Treg proportions in glioma patients (female n=13; male n=14), dataset GSE163108 (27) was used for T cell subtype annotation, and ratios of Treg to CD4^+^ T or total T cells were determined and grouped by genders. Two-sided Mann-Whitney U test implemented by scipy (v. 1.17.1) was performed for statistical analysis, and a p-value < 0.05 was considered statistically significant (35).

### Statistical analysis

Statistical analyses were performed using GraphPad Prism 9. The unpaired two-tailed t test was used for comparison between two groups, whereas one-way or two-way ANOVA was used for multigroup comparisons.

## Results

### DRP-104-resistant tumors become highly susceptible to enzalutamide in immuno-competent hosts

Our group and others have demonstrated potent anti-tumor effects of a glutamine antagonist prodrug, DRP-104, also known as sirpiglenastat, in solid cancers including PCa via direct, cancer cell-autonomous suppression and reprogramming the host’s immune microenvironment (12, 13, 36–39). However, like other targeted therapies, cancer cells treatment can ultimately develop resistance (adaptation) to DRP-104, and we showed that tumor cells can employ multiple intracellular processes to adapt to and become resistant to this pro-drug (13). In the current study, we studied the molecular mechanisms of tumor cells resistance to DRP-104 and identify novel strategies for treating the resistant tumors in an immunologically competent context.

We generated tumors in C57BL/6 host mice from the mouse PCa line TrampC2, following the recently described procedure(13), and devised a treatment/monitoring regimen such that the tumors initially responded to the treatment but ultimately became resistant and progressed in the pilot trial (**Supplementary Fig. 1A**). We decided to use this small cohort of DRP-104-resistant tumors to test their response to the AR antagonist enzalutamide, based on the following rationale. First, AR-targeting treatments cause severe side effects and lower patients’ quality of life (40–45). A better treatment plan may be to use a less toxic agent such as DRP-104 first and use AR-targeted therapies only after the resistance has happened. Second, while the roles of AR signaling on T lymphocytes suggest AR antagonists can be leveraged to enhance anti-tumor immunity(2–4), enzalutamide caused increased immunosuppression in recurrent PCa(7, 8), and its combination with checkpoint blockade has failed to yield clinical benefits(9). Finally, an overwhelming majority of studies involving pre-clinical PCa models and enzalutamide were performed in immunodeficient mice; and in a few cases of studies in immune-competent hosts, TrampC2-derived tumors displayed inconsistent responsiveness to enzalutamide (46, 47). Therefore, we have focused our study on how enzalutamide may interact with tumor microenvironment in immune-competent hosts.

We subjected the small cohort of resistant tumors to enzalutamide monotherapy. Intriguingly, compared to what was found in the prior study(47), even though these DRP-104 resistant tumors were larger (∼300-400 mm^3^ versus ∼50-100 mm^3^), and were treated with less intensity (daily 10 mg/kg versus 15 mg/kg), they displayed significant responses to enzalutamide (**Supplementary Fig. 1B**). Although it was not possible to confidently compare the tumors’ response to therapies across different studies, results from this pilot trial led us to test the hypothesis that, compared to naïve tumors, DRP-104-treated/resistant tumors are more sensitive to enzalutamide in immuno-competent hosts.

Motivated by the above results, we generated a cohort of TrampC2-derived tumors that were not treated with the pro-drug (“naïve” tumors) and a cohort of DRP-104-treated tumors following the protocol described above (“DRP-104-resistant” tumors) (**Supplementary Fig. 1C**). Tumors in both groups, ∼200 mm^3^ each, were treated with vehicle control or with enzalutamide monotherapy of identical regimen (daily 10 mg/kg) (**Supplementary Fig. 1C**). We found that while the naïve tumors showed minimal response to enzalutamide (**Fig. 1A-B**), the resistant tumors were significantly more sensitive (**Fig. 1C-D**). In agreement with the hypothesis that the increased sensitivity was mostly due to the in vivo microenvironment, we found that TrampC2 cells pre-treated with DRP-104 actually became measurably less sensitive to enzalutamide (**Fig. 1E**). The inability of DRP-104 to confer PCa cell’s sensitivity to enzalutamide in vitro was also confirmed in the human PCa cell line, LNCaP (**Supplementary Fig. 1D**), consistent with previous findings that DRP-104 treatment does not alter AR signaling(13). Collectively, these results suggest that DRP-104-resistant tumors become highly susceptible to enzalutamide in immuno-competent host mice.

**Fig. 1.**
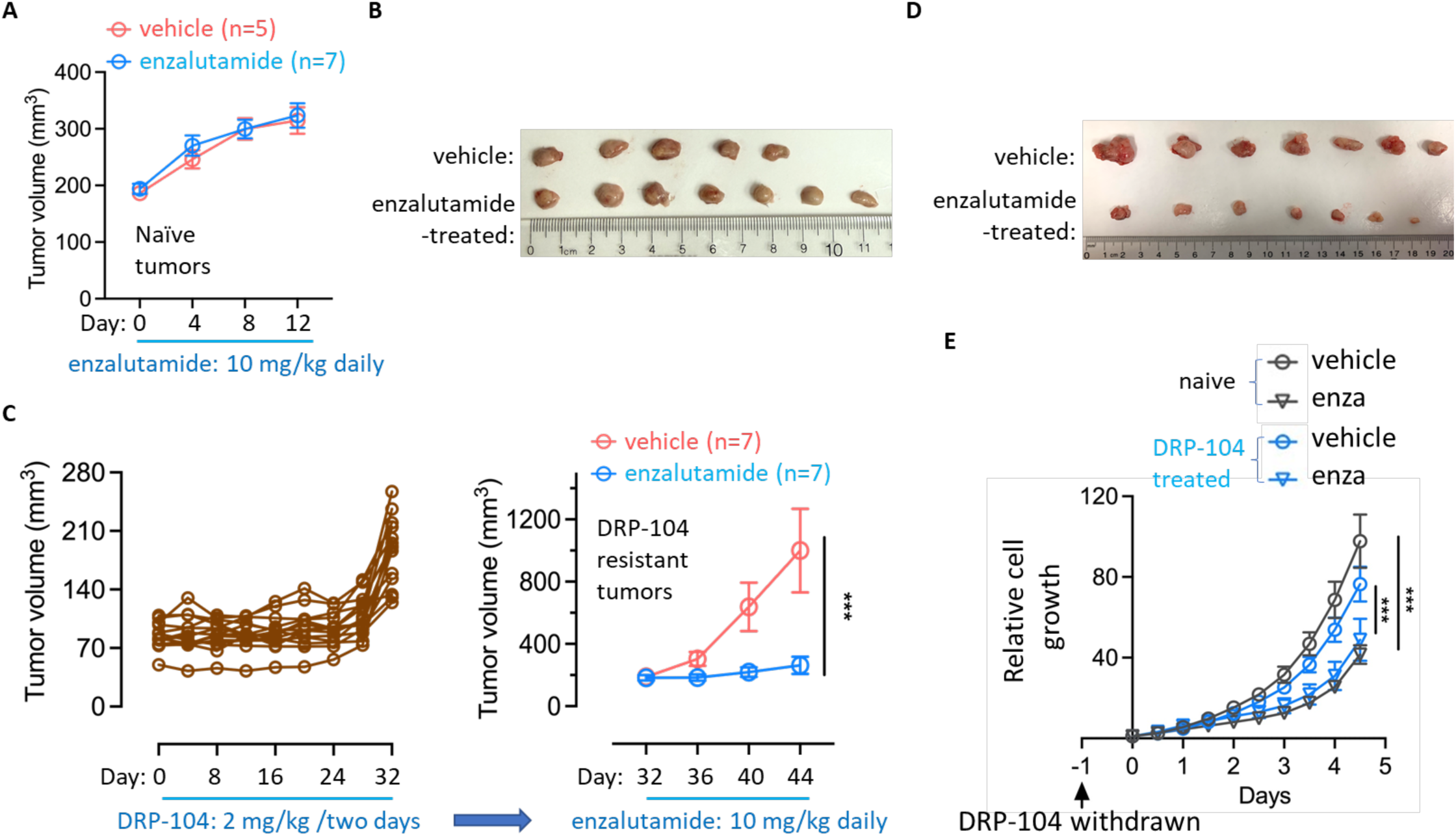
DRP-104-resistant PCa becomes highly susceptible to enzalutamide in immuno-competent host mice. **(A-B)** TrampC2 tumors were generated in C57BL/6 host mice and treated with enzalutamide when their sizes reached ∼200 mm^3^. Tumor volumes were determined (A) and images taken (B). **(C-D)** DRP-104-resistant TrampC2 tumors were generated following the indicated regimen, and when the average tumor volumes reached ∼200 mm^3^, DRP-104 was withdrawn, and enzalutamide monotherapy was initiated. Tumor progression was assessed by tumor volumes (C) and tumor images (D). **(E)** Naïve or DRP-104-treated (2 µM for two weeks) TrampC2 cells were subjected to treatment with enzalutamide (enza), and relative cell propagation was determined via Incucyte. Data was depicted as mean ± SEM. ***p<0.001.

### DRP-104-induced tumor regression and the subsequent tumor resistance are accompanied by altered tumor cell features and immune microenvironment

To gain insights into the tumor’s transitioning from being responsive to becoming resistant to DRP-104, we devised a procedure to generate tumors that were untreated and growing (“Naïve”), tumors that were treated and actively responding (“Responsive”), and tumors that were initially responsive but became resistant and progressing (“Resistant”) (**Supplementary Fig. 2A**). These three groups of tumors were harvested at two time points after the start of the treatment such that the naïve and resistant tumors were actively growing (i.e., a majority of them had sizes of ∼300-600 mm^3^) (**Fig. 2A, B**, **Supplementary Fig. 2B**). Histological analysis of H&E stained slides revealed higher tumor cellularity in the progressing tumors (naïve and resistant groups) compared to the responsive tumors, as expected (**Supplementary Fig. 2C**). Tumors of comparable sizes from the naïve and resistant groups, together with the responsive tumors (n=3 for naïve and resistant groups respectively, n=4 for the responsive group) were subjected to mRNA-seq analysis. Principal component analysis confirmed that these three groups of tumors could be distinguished by the transcriptomic profiles (**Supplementary Fig. 2D**, **Supplementary Table 1**). Further analyses and comparison of the molecular and histological features of these three groups of tumors led to the following findings.

**Fig. 2.**
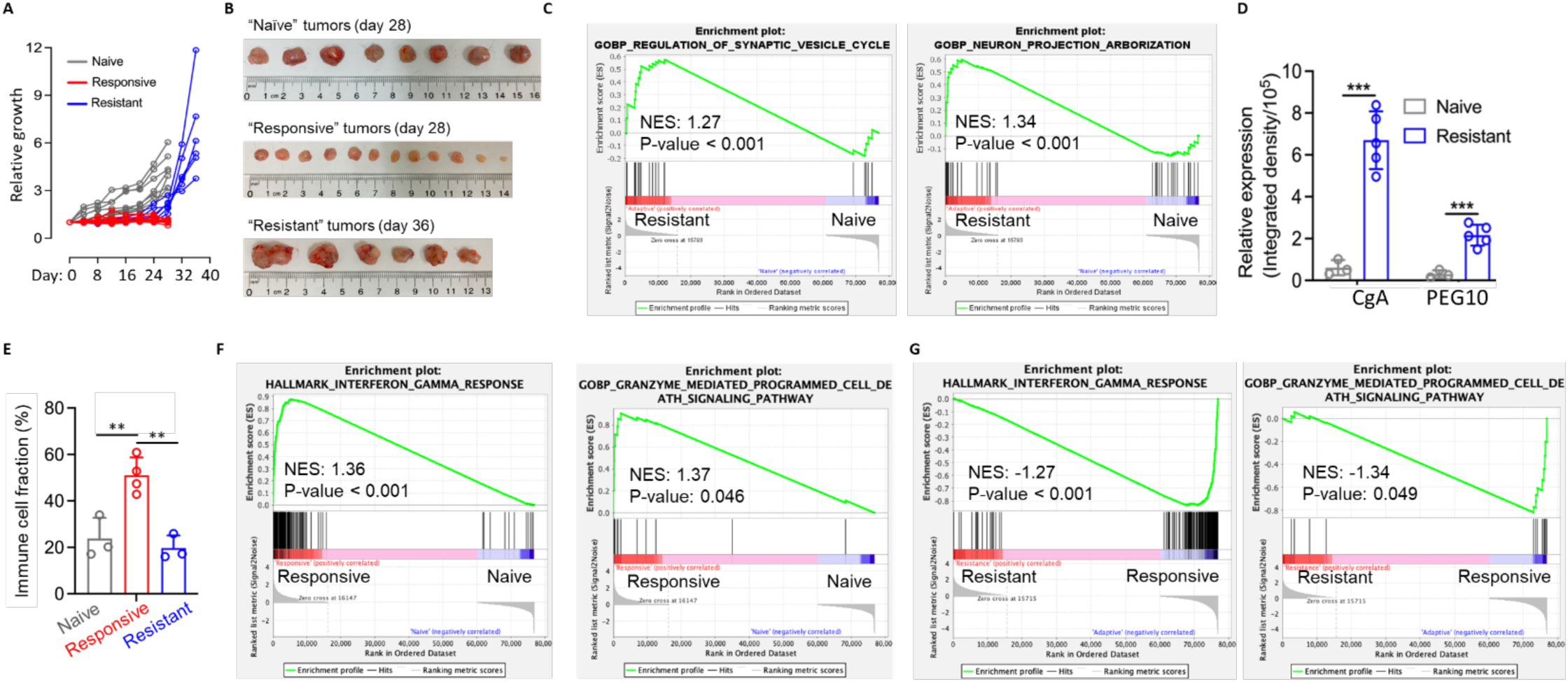
DRP-104-induced tumor regression and the subsequent tumor resistance were accompanied by altered tumor cell features and immune microenvironment. **(A-B)** Three groups of subcutaneous, TrampC2-derived tumors were generated, including those that were untreated (“Naïve”), those that were treated with and responsive to DRP-104 (“Responsive”), and those that were treated with DRP-104, and subsequently became resistant and progressed (“Resistant”). **(C)** The three groups of tumors described in (A-B) were subjected to mRNA-seq and their transcriptomic profiles, and GSEA was performed to reveal distinct features of resistant tumors (versus naïve tumors). **(D)** Naïve and Resistant tumors were used for IHC staining against CgA and PEG10, and their relative IHC signal intensity was shown. **(E)** mRNA-seq data from tumors described in (A-B) were used for cell deconvolution analyses to determine the compositions of the cells in the three groups of tumors. The estimated immune cell fractions of these three groups of tumors were shown. **(F)** mRNA-seq data from tumors described in (A-B) were used for GSEA (gene set enrichment analysis) to compare anti-tumor immunity pathways in the responsive tumors versus the naïve tumors. **(G)** mRNA-seq data from tumors described in (A-B) were used for GSEA to compare anti-tumor immunity pathways in the resistant tumors versus the responsive tumors. **p<0.01. ns: not significant.

First, differential gene and pathway enrichment analysis found that compared to the naïve tumors, the resistant tumors were featured by genes that were associated with neuronal and synaptic activity (**Fig. 2C**). This neuronal feature was confirmed by IHC assays of selected neuroendocrine lineage markers (CgA and PEG10) (**Fig. 2D, Supplementary Fig. 2E**). These results suggest that resistant tumor cells possess molecular features of neuroendocrine identity. To assess whether this resistance-associated neuroendocrine feature was unique to the mouse cell line (TrampC2), we profiled the transcriptomes (mRNA-seq) DRP-104-resistant human cancer cell lines (C4-2 and PC3) developed by us(13) and similarly observed the treatment-induced molecular features of neuroendocrine identity (**Supplementary Fig. 2F**). The detectable change of the tumor cell’s identity is consistent with prior observations that the treated/resistant tumors progressed more rapidly compared to those that were treatment-naïve (**Fig. 1A** and **1C**, **Fig. 2A**), as with cancer therapies in general (in PCa and beyond). Although detailed studies need to be done to further study the molecular mechanisms, we favor the hypothesis that the cell identity switch was likely the result of evolution (adaptation / selection) of tumor cells under the therapeutic pressure(48), instead of a specific, DRP-104-activated transcription of genes associated with the neuroendocrine lineage. Second, cell composition analysis found that compared to the progressing tumors (naïve tumors and resistant ones), responsive tumors were more prominently featured by higher levels of immune cell infiltration (**Fig. 2E**). This finding from deconvolution-based analysis was supported by the observation that most enriched pathways in the responsive tumors were related to immune response and anti-tumor immunity (**Fig. 2F**), and was consistent with the histological features of these tumors described above (**Supplementary Fig. 2C**). The increased immune cell infiltration in the responsive tumors (versus the naïve tumors) was consistent with the previously reported reprogramming of tumor immune microenvironment and anti-tumor immunity induced by the glutamine antagonist pro-drug (13, 37–39). Notably, this trend was reversed in the resistant tumors (**Fig. 2F, 2G**), implicating inefficacy of anti-tumor immunity and/or restored immune suppression in the resistant tumors. Collectively, these results suggest that DRP-104-induced tumor regression and the subsequent tumor resistance were accompanied by alterations in tumor cells’ lineage features as well as in the tumor immune microenvironment.

### Persistent presence of Treg in DRP-104-responsive and resistant tumors

We examined the aforementioned transcriptomic profiles to assess the composition of immune cells in the three groups of tumors. Cell type deconvolution analysis revealed that the fractions of the major immune cells varied greatly across cell types, yet maintained relatively constant among the three groups of tumors with the notable exception of neutrophils and T lymphocytes (**Fig. 3A, Supplementary Fig. 3A**). The functional consequence of the relatively low fraction of neutrophils in the resistant tumors (**Supplementary Fig. 3A**) was unclear, as the roles of neutrophils in cancer progression and immunotherapies are complicated and may be context-dependent (i.e. they can be anti-tumorigenic or immunosuppressive pending on their functional states) (49–52). More significant were the changes of the two cell types that are directly relevant to immunotherapies and immunosuppression, CD8^+^ T cells and Treg: CD8^+^ T cells were readily detectable in only responsive tumors, while Treg became more abundant in both responsive and resistant tumors compared to the naïve ones (**Fig. 3A-B**). These findings from the transcriptomic profile-based analyses were confirmed by IHC staining against CD8 and FOXP3 in the tumors (**Fig. 3C-D**). Subsequently, IF staining against CD4 and FOXP3 in the responsive and resistant tumors confirmed the presence of CD4^+^/FOXP^+^ cells in these tumors (**Fig. 3E**). We used the transcriptomic profiles from aforementioned responsive versus naïve tumors for GSEA (Gene Set Enrichment Analysis(16)), and found the former indeed displayed activated tryptophan metabolism pathway, which broadly impacts immune cells, including Treg, to exert immunosuppression and restrict anti-tumor immunity(53) (**Fig. 3F**).

**Fig. 3.**
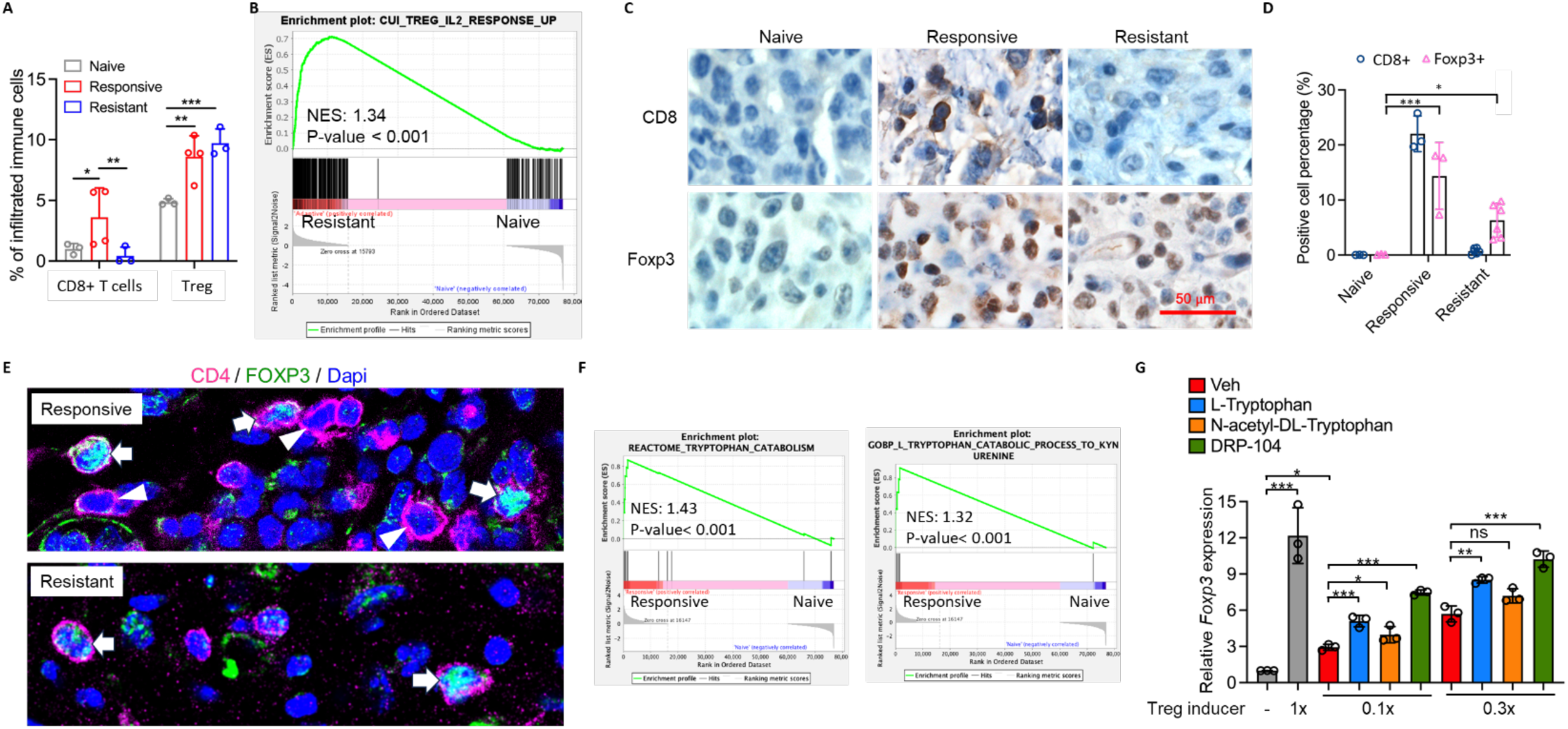
DRP-104 treatment induces transient and persistent presence of CD8^+^ and Treg, respectively, in PCa tumors. **(A-B)** mRNA-seq data from tumors described in Fig. 2 (A-B) were used for cell deconvolution analyses to determine the compositions of CD8^+^ T cells and Tregs in the three groups of tumors (B), and for GSEA to determine the presence of Tregs in tumors (C)**. (C-D)** The naïve, responsive, and resistant tumors were used for IHC assays to determine the relative abundance of CD8^+^ and FOXP3^+^ in the tumors. Representative images (C) and quantifications (D) were shown. **(E)** The responsive and resistant tumors were used for IF staining to confirm the presence of CD4^+^/FOXP3^+^ in these tumors. White arrows denote CD4^+^/FOXP3^+^ Treg cells, and white arrow heads denote CD4^+^ T cells. **(F)** mRNA-seq data from tumors described in Fig. 2 (A-B) were used for GSEA to assess tryptophan metabolism pathways in the responsive tumors (versus the naïve tumors). **(G)** CD4^+^ mouse T lymphocytes obtained via negative enrichment from C57BL/6 mice were used for in vitro Treg differentiation assays with the presence of indicated stimulators/metabolites. The relative expression of Foxp3 was determined by RT-qPCR as a way to estimate the efficiency of Treg differentiation. Data were depicted as mean ± SEM. *p<0.05, **p<0.01, ***p<0.001. ****p<0.0001, ns: not significant.

Tregs enable tumors’ evasion of immune surveillance in tumorigenesis and impair the antitumor effector cells (e.g., CD8^+^ T cells) to limit the efficacy of immunotherapies, making them a crucial therapeutic target(54–56). Elevated infiltration and expansion of Tregs in tumors commonly occur in response to various types of cancer immunotherapies such as immune checkpoint blockade, CAR T, and cancer vaccines (57–59). Thus, the above results suggest that such therapy-associated immunosuppression also occurs after DRP-104-mediated glutamine/purine blockade treatment.

While the presence of Treg could simply be the host immune system’s natural response to therapeutically stimulated anti-tumor immunity, several independent lines of evidence suggest that, while serving as a potent immunotherapeutic agent, the glutamine antagonist prodrug likely also contributed to Treg’s differentiation and/or functions in tumors. DRP-104 has an acetylated tryptophan moiety, which is known to be immunosuppressive upon the TDO2/IDO1-mediated catabolism (38, 53). Prior metabolite profiling results showed that human PCa cells treated with DRP-104 had significantly higher levels of acetyl-tryptophan and activated tryptophan metabolism in tumor cells (13). Additionally, re-examination of metabolite profiling data from PCa cell lines(13) treated with the pro-drug revealed that the treatment also resulted in high levels of metabolites that can serve as the ligands for the aryl hydrocarbon receptor (AhR)(60, 61): the nuclear receptor that is essential to Treg development (61, 62) (e.g., indoleacrylic acid; **Supplementary Fig. 3B**). Although such DRP-104-induced metabolites were initially observed in tumor cells(13), it was conceivable that they were also present in microenvironment and contributed to a more immunosuppressive landscape in the treated (responsive and resistant) tumors. Finally, we assessed the direct effects of DRP-104 on CD4^+^ T cells. Using in vitro Treg differentiation assays and the expression of *Foxp3* as the readout, we found that DRP-104, similar to tryptophan and acetyl tryptophan, augmented the expression of *Foxp3* (induced by low doses of a Treg inducing mixture of IL-2, TGFβ, and all-trans retinoic acid (ATRA)) (**Fig. 3G**). Although the mechanism underlying this direct effect of DRP-104 on *Foxp3* and its relevance to the in vivo context remain to be elucidated, this result points to DRP-104’s direct effects on Treg’s differentiation.

Collectively, findings from in vivo and in vitro models support that, similar to other immunotherapeutic strategies (57–59), the anti-tumor immunity induced by the glutamine/purine-blockade treatment is accompanied by immunosuppressive effects (e.g., by promoting Treg) that may limit its long-term therapeutic efficacy.

### Combining enzalutamide and DRP-104 depletes Treg from tumors and results in superior therapeutic efficacy

The DRP-104-induced changes in the CD8^+^ T cells and Treg and DRP-104 resistant tumors’ superior response to enzalutamide led us to examine the profiles of these T cells in the tumors (DRP-104- naïve or resistant) that were treated with enzalutamide (or with vehicle control, as depicted in **Supplementary Fig. 1C**). We took the tumors that were harvested at the end point of the treatments (tumor images were shown in **Fig. 1B** and **1D**), and performed IHC staining against CD8 and FOXP3. CD8^+^ or FOXP^+^ cells were barely detectable in both vehicle and enzalutamide-treated tumors that were DRP-104-naïve (**Fig. 4A** and **Supplementary Fig. 4A**). This was not surprising, as the enzalutamide regimen used in the trial only minimally affected these tumors (as shown in **Fig. 1A**). In contrast, tumors that were previously treated with and became resistant to DRP-104 were infiltrated with readily detectable CD8^+^ T cells upon enzalutamide treatment (**Fig. 4A** and **Supplementary Fig. 4A**), suggesting enzalutamide promoted the presence of CD8^+^ T cells that were likely initially induced by DRP-104. While these analyses did not assess the effects of enzalutamide on the anti-tumor immunity of these T cells, the results were consistent with the potent therapeutic efficacy observed (as shown in **Fig. 1C**) and supported the conclusion that enzalutamide augments CD8^+^ T cells’ anti-tumor activity(2).

**Fig. 4.**
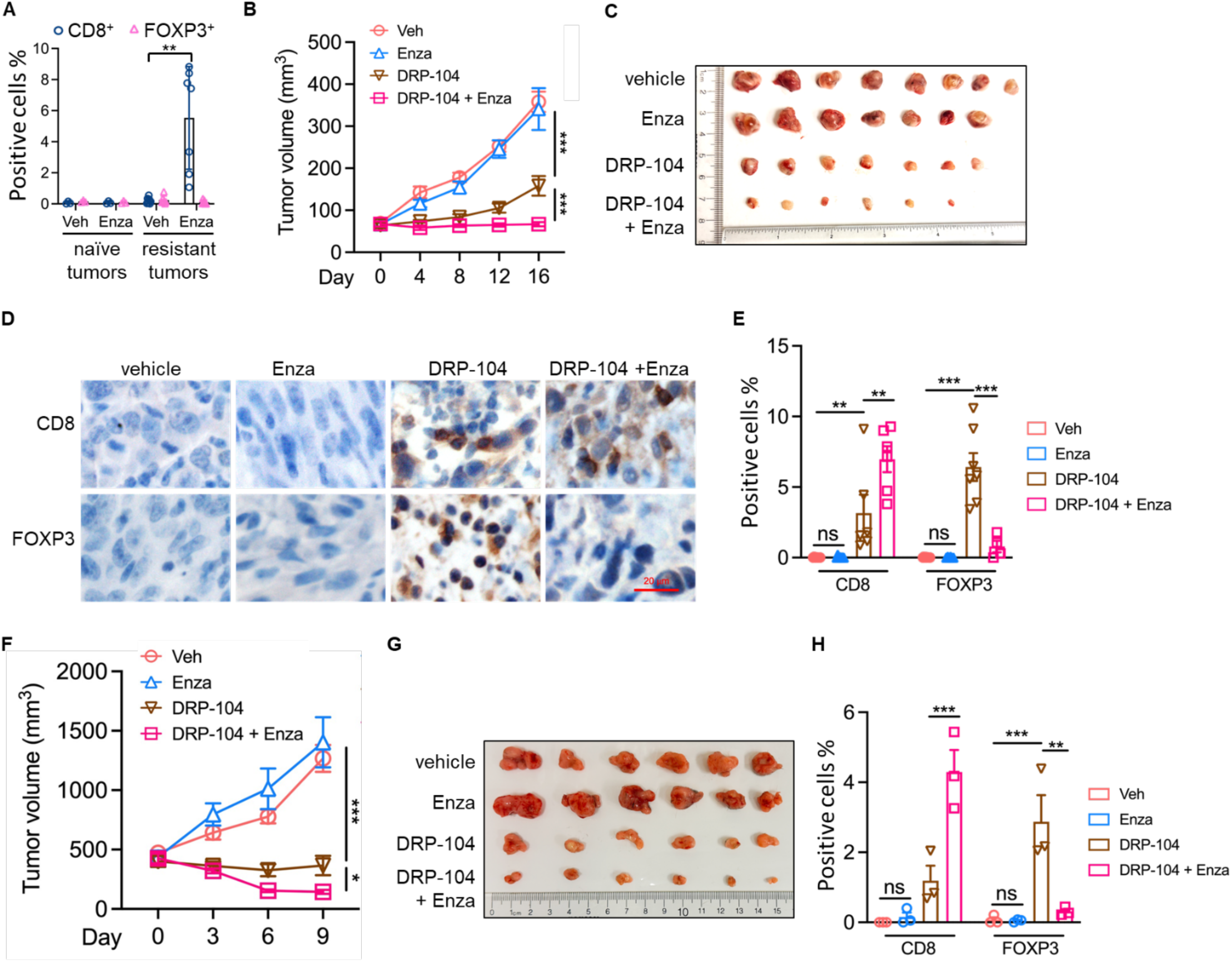
Combining DRP-104 and enzalutamide results in superior therapeutic efficacy and depletes Treg from tumors. **(A)** DRP-104 naïve or resistant tumors were treated with vehicle control or with enzalutamide for 12 days, at which point most vehicle-treated DRP-104-resistant tumors reached the humane endpoint (tumor volumes of ∼1000 mm^3^), and all tumors were harvested for IHC analysis. The quantification results were shown (Veh: vehicle; Enza: enzalutamide). **(B-C)** C57BL/6 mice bearing subcutaneous tumors derived from TrampC2 were used for four arms of treatments: vehicle control, Enza monotherapy (10 mg/kg) daily, DRP-104 (2 mg/kg) every two days, or both agents, and the tumor growth was determined. The growth curves (B) and images of tumors harvested at the human endpoint (C) were shown. **(D-E)** Tumors shown in (C) were used for anti-CD8 and anti-FOXP3 IHC staining, and the representative IHC images (D) and the quantification of the positively stained cells (E) were shown. **(F-G)** C57BL/6 mice bearing subcutaneous tumors derived from PPR, a mouse NEPC-like cell line, were used for four arms of daily treatments: vehicle control, Enza monotherapy (10 mg/kg) daily, DRP-104 (2 mg/kg) every two days, or both agents, and the tumor growth was determined. The growth curves (F) and images of tumors harvested at the human endpoint (G) were shown. **(H)** Tumors shown in (G) were used for anti-CD8 and anti-FOXP3 IHC staining, and the quantification of the positively stained cells was shown.

Notably, no Treg was detected in the enzalutamide-treated DRP-104-resistant tumors (**Fig. 4A**). As their vehicle-treated counterparts (harvested at the humane endpoint) had no detectable Treg to start with, it was not possible to assess any suppressive effects of enzalutamide on Treg in this cohort of tumors. This prompted us to test a combination therapy strategy, i.e., generating TrampC2-derived tumors and treating them with DRP-104 or enzalutamide as monotherapy therapies or applying both agents simultaneously. Importantly, we observed that while enzalutamide or DRP-104 as a monotherapy had no effects or effective suppression on the tumors, combining both agents led to superior tumor suppression (**Fig. 4B, 4C**). IHC staining against CD8 and FOXP3 on these tumors revealed that while enzalutamide monotherapy did not induce CD8^+^ T cells in the tumors, it augmented the presence of CD8^+^ T cells when used in combination with DRP-104 (**Fig. 4D-E**). Most remarkably, when compared to tumors treated with DRP-104 monotherapy, the tumors treated with the combination therapy had significantly lower levels of FOXP3^+^ cells (**Fig. 4D-E**), suggesting that enzalutamide effectively mitigated tumor-infiltrated Treg induced by DRP-104 therapy.

The enzalutamide-mediated depletion of Treg from the DRP-104-treated tumors raised two questions. The first was whether this was simply a TrampC2 model-specific effect. Additionally, even though we have observed that the DRP-104 naïve TrampC2 tumors had minimal susceptibility to the enzalutamide regimen used and that the DRP-104-resistant tumors displayed neuroendocrine-like molecular features, it was possible that the effects of enzalutamide were specific to adenocarcinoma (i.e., AR-driven PCa) but not applicable to tumors of other lineages. To address these questions, we generated tumors derived from an extremely aggressive, neuroendocrine PCa (NEPC)-like tumor cell line, which was derived from a mouse model of *Tp53*/*Pten*/*Rb1* deficiency without AR expression in tumor cell (14). Our first trial testing the monotherapies and the DRP-104 plus enzalutamide combination therapy found that using the standard regimens (2 mg/kg for DRP-104 every two days and 10 mg/kg for enzalutamide every day, respectively, as monotherapy or in combination), enzalutamide had no effects on these tumors as expected (**Supplementary Fig. 4B, 4C**). However, DRP-104 displayed strong suppressive effects as a monotherapy, while the combination therapy showed more effective tumor suppression after three treatments (day 6, at which point the trial reached the humane endpoint due to the very aggressive growth of the vehicle and enzalutamide-treated tumors) (**Supplementary Fig. 4B, 4C**). These results prompted us to perform a second trial in which the tumors were treated more intensely (i.e., from every two days to daily treatments of DRP-104, with the same drug doses), such that the trial’s humane endpoint was not reached until day 9. This second trial confirmed the superior therapeutic efficacy of the combination therapy (**Fig. 4F-G**). IHC assays on the treated tumors again revealed enzalutamide-augmented infiltration of CD8^+^ T cells, and most remarkably, the near complete depletion of FOXP3^+^ cells from the tumors (**Fig. 4H**, **Supplementary Fig. 4D**). Together, these results from two in vivo tumor models suggest that enzalutamide potently promoted CD8^+^ T cells and most importantly, depleted Treg from the tumors in the context of DRP-104-mediated glutamine/purine blockade therapy.

### Enzalutamide suppresses Treg differentiation and disrupts the interaction of AR and AhR in the nucleus of mouse CD4^+^ T cells

The PCa cell-autonomous oncogenic effects of the AR signaling have been the focus of most studies in the field of PCa research. Interestingly, recent research has uncovered promoting effects of AR signaling on CD8^+^ T cell exhaustion(2). This, together with the well-established immunosuppressive property of Treg in tumorigenesis and in cancer immunotherapies in the context of PCa bone metastases(63, 64), prompted us to test whether enzalutamide directly mitigates Treg development. We performed in vitro experiments using mouse spleen-derived CD4^+^ T cells for Treg differentiation assays, using tumor cell-conditioned, DRP-104^+^ culture media supplemented with exogenous IL2 and TGFβ (plus ATRA in some cases). We found that the tumor cell-conditioned media effectively induced *Foxp3* expression, which was attenuated by enzalutamide (**Fig. 5A-B**). This inhibitory effect of enzalutamide on Treg differentiation was also observed when using an optimized Treg inducer master mixture (ImmunoCult from Stemcell Tech, cat^#^ 10957) for Treg differentiation (**Fig. 5C**).

**Fig. 5.**
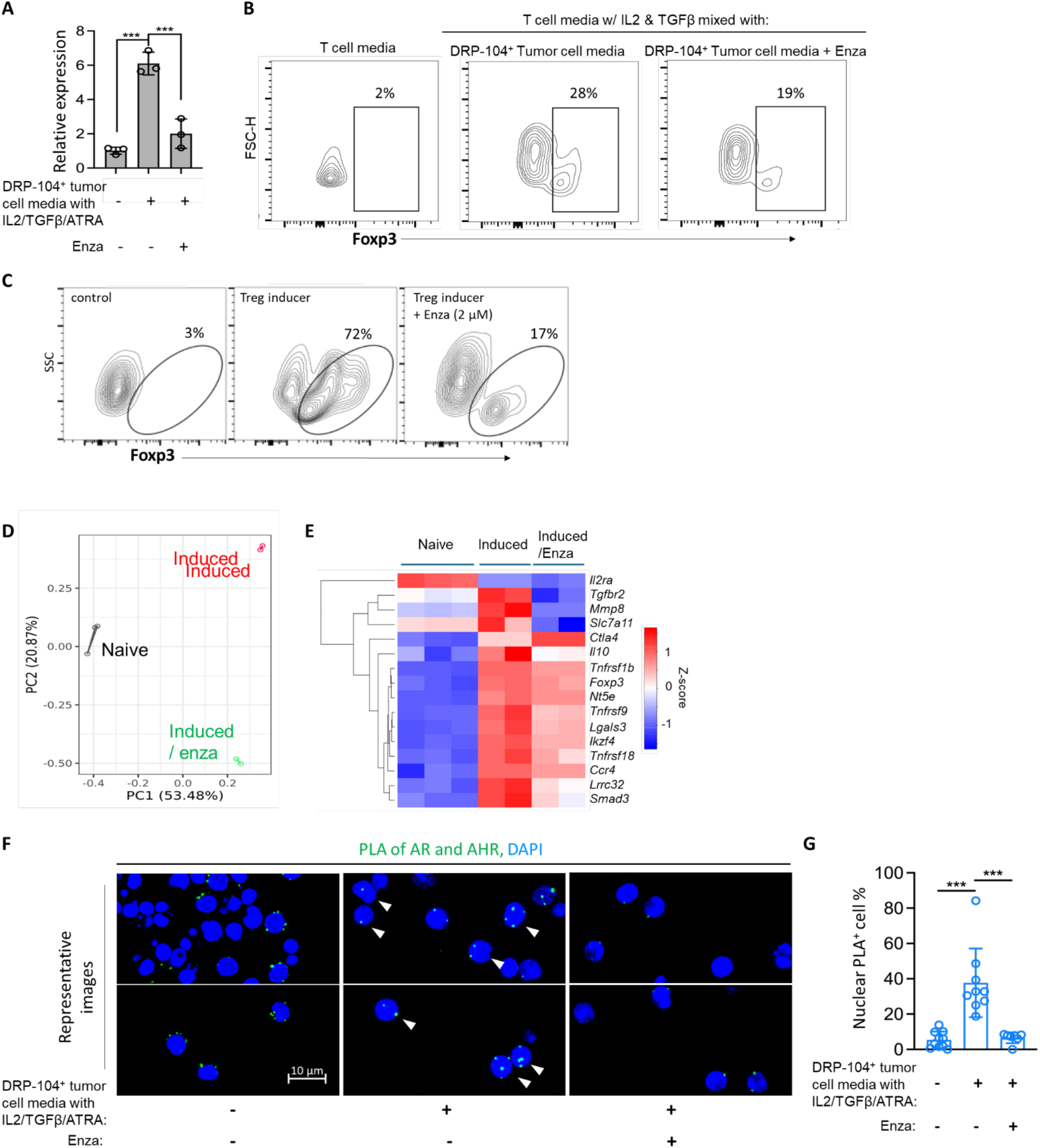
Enzalutamide attenuates Treg differentiation and disrupts the AR and AhR complex in the nucleus. **(A-B)** CD4^+^ T cells were obtained from C57BL/6 mouse spleens and in vitro Treg differentiation using indicated conditional media, and Treg differentiation was assessed by (A) RT-qPCR analysis of *Foxp3* expression and (B) intracellular anti-Foxp3 staining and FACS analysis. **(C)** CD4^+^ T cells were subjected to Treg differentiation by using a commercial, optimized Treg inducer mixture, and intracellular anti-Foxp3 staining and FACS were performed to assess Treg differentiation. **(D)** Principal component analysis for the transcriptomic profiles from the three populations of cells. **(E)** A heat map showing the changes in the expression of 16 genes associated with Treg differentiation. ***p<0.001. Enza: enzalutamide. **(F-G)** CD4^+^ T cells from mouse spleens were used for Treg differentiation, without or with enzalutamide, and PLA was performed. **(F)** Representative images (two images from each condition) were shown. Note the PLA signal (arrowed) in the nucleus in the presence of Treg inducer (middle panel). **(G)** Quantification results (∼10 fields from each group presented as one dot in the plot, including n=1130, 1436, and 1843 cells, respectively, for the three groups, were quantified). *p<0.05, **p<0.01, ***p<0.001.

To further assess the effects of enzalutamide on Treg differentiation, we enriched CD4^+^ T cells from mouse splenocytes via negative selection. We then subjected them to Treg differentiation with or without enzalutamide to obtain three populations: Naïve cells (CD4^+^-enriched cells), Induced cells (i.e., CD4^+^-enriched cells subjected to Treg induction), and Induced/enza cells (i.e., CD4^+^-enriched cells subjected to Treg induction in the presence of enzalutamide). Total RNAs were prepared from these populations. Due to the extremely low amount of total RNA obtained from these populations, non-ribosomal RNA sequencing, instead of typical mRNA-seq, was used to profile the effects of enzalutamide on the transcriptomes. As a result of the low abundance of non-ribosomal RNA obtained, only seven libraries, including three, two and two replicates from the three populations of cells respectively, were successfully prepared and sequenced for downstream analyses. Despite this limitation, principal component analysis revealed enzalutamide’s profound effects on the induced cells (**Fig. 5D, Supplementary Fig. 5A**). We identified differentially expressed genes (DEGs) in the Induced cells (versus the Naïve cells), and selected those whose expression was affected by enzalutamide (i.e., DEGs in the Induced/enza cells versus Induced cells). We then used them (n=1392 genes) to estimate the immune cell types/states via CIEC(24). This analysis revealed that enzalutamide broadly affected the expression of genes associated with multiple lineages of immune cells including T, B, and myeloid cells (**Supplementary Fig. 5B**). We then specifically examined a set of genes associated with the differentiation/stability or the immune suppressive function of Treg and found that, with the exception of *Ctla-4,* the induced expression of these genes was mitigated by enzalutamide (**Fig. 5E**).

The mitigating effects of enzalutamide on Treg prompted us to test whether AR was involved in Treg differentiation. Previous studies found that in certain PCa and breast cancer cell lines, upon engagement by its ligand (3-methylcholanthrene, 3MC), AhR could form complexes with AR with unclear pathological consequences (65) or causing the latter’s degradation(66). We therefore assessed whether AR and AhR interacted in the CD4^+^ T cells by proximity ligation assays. The assays confirmed AR and AhR interaction in the cytosol of CD4^+^ T cells, and the interacting proteins relocated to the nucleus under the Treg-inducing condition, which was significantly diminished when enzalutamide was present (**Fig. 5F-G**). These results suggest that AR interacts with AhR directly or indirectly and work together to drive Treg differentiation and/or stabilization, and it likely does so via functionally cooperating with the AhR.

### Gene expression analysis implicates AR signal in shaping the Treg landscape in cancers

Several previous studies in the field of autoimmune diseases have linked AR to Treg biogenesis and/or functions. Using *Ar*-knockout mouse models, it was found that AR signal promotes Treg’s immune suppressive function during allergic airway inflammation(67), and that *Ar* loss resulted in fewer thymic and peripheral Treg cells in male mice(68). In a rat model of experimental autoimmune orchitis, testosterone was found to stimulate Treg expansion(69). AR signaling affects Treg cells in complicated gender and age-dependent manners in in vitro analyses of human PMBC. While an AR ligand can stimulate *Foxp3* expression, female PBMC were found to have higher fractions of Treg (CD4^+^CD25^+^Foxp3^+^) cells(70). Additionally, it was found that, in general, male PBMCs had higher *Foxp3* message mRNA levels than female PBMCs(71). Together, these findings suggest a discordance between Treg biogenesis in response to AR signal, Treg homeostasis, and the inherent complexity presented by assays used. Nevertheless, several independent lines of evidence suggest AR’s Treg-promoting effects in healthy human PBMCs and in cancers. By re-analyzing the available gene expression data from bulk mRNA-seq, DICE(26), we found that among immune cells from human PMBCs, memory (effector) Treg displayed the highest level of *AR* expression (**Fig. 6A**). Furthermore, Treg had higher levels of *AR* expression than naïve CD4^+^ T cells regardless of the gender (**Fig. 6B**). To assess the potential involvement of AR signaling in Treg biogenesis/homeostasis in cancers, we utilized the available scRNA-seq datasets from human cancers in which a large number of CD8^+^ and CD4^+^ T cells were identified: GSE150430 (nasopharyngeal carcinoma), GSE139324 (head and neck cancer), and GSM4972161 (glioma) (27–29). We first examined the relationship between AR signaling activity, defined by Hallmark’s androgen response gene set(30), and CD8^+^ T cell exhaustion score, defined by the expression of the previously established exhausted CD8^+^ T cell gene panel (33). This analysis revealed the trend of positive correlations between the AR activity and the exhaustion scores in CD8^+^ T cells in the tumors (**Fig. 6C**), which was in agreement with the established roles of AR signaling in driving CD8^+^ T cell exhaustion in cancers (2, 3). Intriguingly, a similar trend of positive correlations was also observed between AR activity and Treg scores (defined by the expression of four Treg-associated marker genes: *FOXP3*, *CTLA-4*, *CCR8* and *TNFRSF9*(31, 32)) in CD4^+^ T cells in the tumors (**Fig. 6D**). Finally, the established link between gender and immune landscapes in the glioma (5, 72, 73) prompted us to examine the relative Treg abundance in female and male patients’ (n=13 and 14, respectively) gliomas. This analysis revealed higher proportions of Treg among CD4^+^ T cells or total T cells in the latter group (**Fig. 6E**). Collectively, these results support our observation in experimental models and implicate AR signaling as an important factor in shaping the Treg landscape and immunosuppression in tumors. They also add to the previous finding that AR signal was generally correlated with an immunologically cold tumor microenvironment (74, 75).

**Fig. 6:**
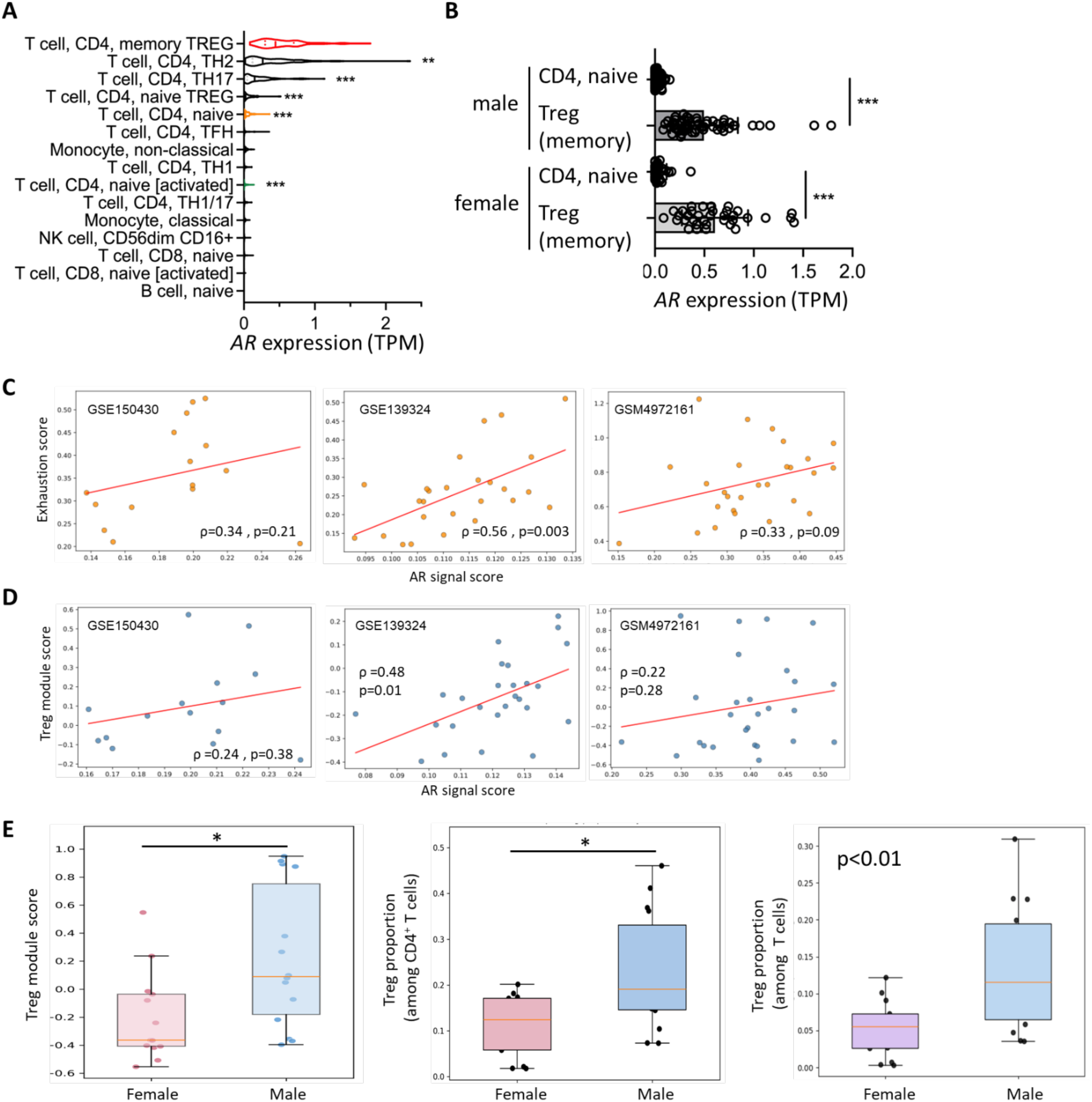
Gene expression analysis implicates AR signal in shaping the Treg landscape in human PBMCs and in tumors. **(A-B)** AR expression (defined by TPM) in various immune cell types in human PBMC (data from https://dice-database.org/). In (A), p values denote the comparison between the other cell types versus memory Treg. **(C-D)** Three scRNA-seq datasets were used to assess the correlations between AR signal activity and T cell exhaustion scores in CD8^+^ T cells (shown in C) or Treg module scores in CD4^+^ T cells (shown in D) in human tumors (nasopharyngeal carcinoma, head and neck cancer, and glioma; each spot represented one tumor). Spearman correlation coefficient (ρ) and p values were shown. **(E)** The glioma scRNA-seq dataset (GSM4972161) was used to compare the relative Treg abundance in tumors, as measured by Treg module scores and the proportions of Treg among CD4^+^ T cells or among total T cells, in female versus male patients. *p<0.05, ***p<0.001.

### CD4^+^ T cells exposed to enzalutamide during Treg induction display diminished suppressive effects on CD8^+^ T cell-mediated tumor cell killing

The above findings prompted us to assess the effects of enzalutamide on Treg’s immunosuppressive activity. We have developed an anti-Gpc3 peptide vaccine strategy on the basis that it would not only serve this purpose but also add to the repository of peptide vaccines for studying cancer immunotherapies. Gpc3 was chosen as this oncofetal protein is specifically present in subsets of many types of cancers, including PCa(76), lung cancers(77), and brain tumors (including glioma)(78, 79), and peptide-based vaccination targeting Gpc3 has been investigated for treating liver cancer(80, 81). We have tested the immunogenicity of nine Gpc3-derived peptides predicted to be presented by MHC-I/II in the C57BL/6 mouse strain by The Immune Epitope Database (IEDB) (**Supplementary Fig. 6A**). We carried out a vaccination trial in which mice were vaccinated twice (on day 1 and day 25) with candidate peptides in the presence of two adjuvants (Addavax + CpG 1826), and splenocytes from the vaccinated mice were harvested (on day 39) for peptide-specific IFNɣ Enzyme- Linked ImmunoSpot (ELISPot) assays. This screening has identified three novel immunogenic peptides (peptide ^#^1 for MHC-II and ^#^5 & ^#^7 for MHC-I) (**Supplementary Fig. 6B-C**). A second vaccination trial using the mixture of these three peptides confirmed the immunogenicity of this triple-peptide mixture (**Fig. 7A**). Subsequently, CD8^+^ T cells isolated from the splenocytes of the triple peptide-vaccinated mice were re-challenged with the triple peptide mixture and used to validate the Gpc3-dependent cytotoxic killing. As expected, these activated CD8^+^ T cells did not eliminate the Gpc3-negative parental TrampC2 cell line (**Fig. 7B, Supplementary Fig. 6D**), yet they potently eradicated tumor cells expressing exogenous Gpc3 (**Fig. 7C, Supplementary Fig. 6D**). This cytotoxic killing of tumor cells was strongly suppressed by CD4^+^ T cells that were subjected to Treg differentiation, but less so when the differentiation was induced in the presence of enzalutamide (**Fig. 7C, Supplementary Fig. 6E**). These results support the in vitro Treg differentiation assays and further implicate AR signaling in promoting Treg’s differentiation and/or immunosuppressive functions and nominate enzalutamide as a potentiating agent for immunotherapies.

**Fig. 7.**
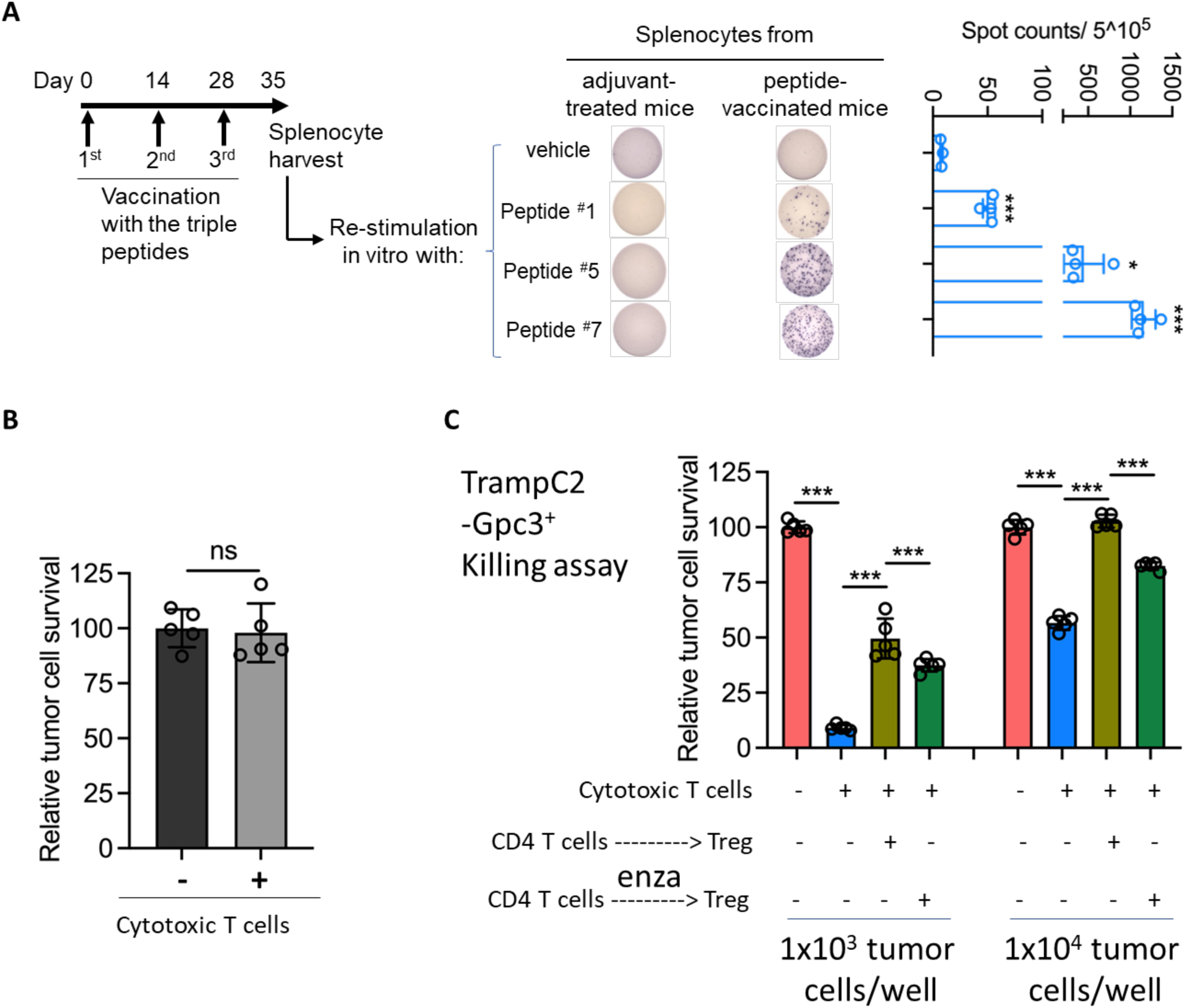
CD4^+^ T cells exposed to enzalutamide during Treg induction displayed diminished suppressive effects on CD8^+^ T cell-mediated tumor cell killing. **(A)** Splenocytes from mice vaccinated with triple peptide mixture (^#^1, 5, and 7) were used for in vitro stimulation with PBS or each indicated peptide, and anti-IFNɣ ELISPot assay was performed. Quantification and representative images were shown. **(B-C)** CD8^+^ cytotoxic T cells enriched from splenocytes from triple peptide-vaccinated mice were used for in vitro CTL killing assays using **(B)** parental TrampC2 cells or **(C)** exogenous Gpc3 expression TrampC2 (Gpc3^+^) cells. In (C), CD4^+^ T cells that were subjected to Treg differentiation with or without the presence of enzalutamide (enza) were added to the CTL assay mixtures (CD4^+^ T to CD8^+^ T cells ratio was ∼5:1). Immune cells were subsequently removed, and relative numbers of tumor cells were quantified by CTG assays. In (C), results from two independent experiments (with different numbers of tumor cells used per well) were shown *p<0.05, ***p<0.001, ****p<0.0001.

## Discussion

In this study, we demonstrated that when treated with the glutamine antagonist prodrug DRP-104, PCa tumors initially responded but subsequently became resistant, and the treated tumors, even while being responsive to the drug, were infiltrated with Treg cells. We further showed that the resistant tumors were highly susceptible to enzalutamide, and the superior response was accompanied by Treg depletion from the tumor tissues. Simultaneously treating naïve tumors with DRP-104 and enzalutamide also resulted in similar observations. Subsequent mechanistic studies revealed that enzalutamide suppresses Treg differentiation, which is accompanied by the disruption of AR and AhR interaction in the nucleus of CD4^+^ T cells. Further gene expression analysis using bulk RNA-seq and reanalysis of gene expression datasets from human PBMC and human cancers implicated AR as one important factor in promoting Treg biogenesis and in defining the Treg landscape in human cancers. Finally, in vitro assays using cytotoxic CD8^+^ T cells from an anti-Gpc3 based vaccination model demonstrated that Treg cells induced in the presence of enzalutamide had diminished suppressive effects on CD8^+^ T cell’s cytotoxic killing of tumor cells. Together, these findings uncover a mechanism of PCa’s resistance to DRP-104-mediated immunotherapeutic effects, elucidate a novel link between enzalutamide and Treg in the context of immunotherapies, and inform future efforts to devise efficacious immunotherapeutic strategies, as elaborated below.

While independent studies have demonstrated potent anti-tumorigenic effects of DRP-104 in multiple cancer types by direct tumor cell suppression and anti-tumor immunity(12, 36–39), our recent study has identified a tumor cell-autonomous mechanism of resistance to this pro-drug (i.e., by compensatory intracellular processes that enhance purine supplies) (13). Findings presented in the current study suggest that a tumor cell non-autonomous, microenvironment-driven mechanism, i.e., the therapy-induced infiltration of Treg cells, likely also facilitates tumor’s resistance/recurrence. It was possible that the Treg infiltration, occurring while tumors were actively responsive to the pro-drug, was simply due to various cytokines and chemokines in the tumor microenvironment in response to the robust activation/infiltration of anti-tumorigenic CD8^+^ T cells, reminiscent of what was observed in glioblastoma tumors treated with chimeric antigen receptor (CAT) T cells (58). However, the association of the pro-drug with multiple immunosuppressive metabolites, such as tryptophan, suggests that the pro-drug, while actively stimulating anti-tumor immunity, likely also facilitated Treg’s biogenesis in situ and/or infiltration into tumors. The precise molecular mechanism underlying DRP-104’s stimulating effects on Treg biogenesis remains unclear. It was previously demonstrated that DRP-104, which possesses an N-acetyl-tryptophan moiety, could be processed by serine proteases in tumors and by carboxylesterases in the gastrointestinal tissue(38). Thus, it is possible that DRP-104-derived metabolites such as N-acetyl-tryptophan, which can be converted to Treg-stimulating tryptophan(82), is released by tumor cells or non-tumor cells into the tumor microenvironment and subsequently acts on T cells. Alternatively, as a non-mutually exclusive mechanism, the pro-drug may also be processed directly by and affect T cells, as shown by the results of the in vitro Treg differentiation assays. We are mindful that our assays did not assess the contribution of the N-acetyl tryptophan moiety to the biogenesis of Tregs in the tumor tissues or the Treg differentiation in vitro. Nevertheless, the findings suggest the potential adverse or beneficial effects of the chemical moieties when designing pro-drugs. Furthermore, the potential effects of the non-effector moieties should also be considered when devising combination therapies, as illustrated by our findings: Enzalutamide appears to effectively mitigate Treg’s presence and vastly improve the therapeutic efficacy of DRP-104, serving as an example of turning an otherwise undesired property of the pro-drug into therapeutic benefits.

The observation that DRP-104-resistant tumors displayed neuroendocrine features is particularly notable for two reasons. First, compared to the naïve tumors, the DRP-104-resistant tumors were more neuroendocrine lineage-like, yet they also became highly susceptible to enzalutamide treatment in vivo (but not in vitro). This provides strong evidence that the superior efficacy of enzalutamide against these tumors was due to its microenvironmental effects (i.e., enhancing anti-tumor immunity), rather than through direct suppression of tumor cells. Second, the DRP-104 treatment and the subsequent emergence of neuroendocrine-like features in the resistant tumors resemble treatment-induced NEPC, which most commonly arises after adenocarcinoma receives and becomes resistant to AR-targeting therapies and is thought to be driven by lineage plasticity (83–85). Unlike AR-targeting therapies, DRP-104 primarily exerts its therapeutic efficacy by disrupting glutamine utilization/purine supplies (tumor cell-autonomous effects) and stimulating anti-tumor immunity(12, 13, 36–39). Thus, DRP-104-driven emergence of neuroendocrine features echoes the previous observation that PCa subjected to and surviving radiation therapy also gained features of neuroendocrine PCa(86), suggesting that luminal cancer cells become adaptive to the therapeutic pressures and ultimately neuroendocrine phenotype(85). These findings support our previous observation that neuroendocrine-like cells pre-exist in treatment-naïve adenocarcinoma and are selected by AR-targeting therapies(87). Together, they suggest that the emergence of NEPC-like tumors is an outcome of therapeutic stress-driven selection, adaptation, and evolution of tumors at clonal, oligoclonal, and populational levels. They also highlight the need to prevent the emergence of NEPC and/or treat NEPC using systematic approaches (e.g., immunotherapies), rather than targeting selective pathways in tumor cells.

The finding that enzalutamide mitigates Treg differentiation advances our understanding of the roles of AR signaling in Treg biogenesis and provides a new direction for immunotherapeutic designs. The link of *Ar* and Treg’s function/homeostasis has been described in an *Ar*-knockout allergic airway inflammation mouse model previously(67). Additionally, *Ar* knockout leads to fewer thymic and peripheral Treg cells in male mice(68), and in a rat EAO model, testosterone, an AR ligand, stimulated Treg expansion(69). Our findings described here suggest that AR activity in CD4^+^ T cells also promotes Treg in the context of cancer immune microenvironment and cancer immunotherapies. Notably, AR can form complexes with AhR, particularly in the presence of their ligands, in human PCa and breast cancer cell lines, yet the functional outcomes of their interaction in tumor cells remain unclear (65, 88) (89). Our finding that AR and AhR interact in the nucleus of CD4^+^ T cells undergoing Treg differentiation induction is a novel finding suggesting that AR likely functionally cooperates with AhR in directing/promoting Treg differentiation. Furthermore, the enzalutamide-mediated disruption of the AR-AhR complexes in the nucleus suggests the mitigating effects of enzalutamide on Treg were exerted through targeting AR signaling, rather than through AR-independent off target effects. These findings are complementary to the previously described roles of AR signaling in promoting T cell exhaustion(2). Together, they suggest that the AR signaling in T cells play important roles in shaping immunosuppressive cancer microenvironment as well as contribute to the gender-dependent differences in auto-immunity, in cancers’ immune microenvironment, and in their responses to immunotherapies (3–6, 72).

Our study has important therapeutic implications. First, AR’s connection to CD8^+^ T cell exhaustion and its association with an immunologically “cold” tumor microenvironment in general support leveraging enzalutamide (or potentially other AR antagonists) for improving cancer immunotherapies(2–4, 74, 75). Our findings further strengthen the argument for such an approach, as enzalutamide can exert dual beneficial effects of potentiating the cytotoxic activity of anti-tumorigenic CD8^+^ T cells while simultaneously mitigating the immunosuppressive effects of Treg cells. Of note, in PCa, enzalutamide and other AR-targeting drugs designed to directly suppress AR-dependent tumor cells have been associated with enhanced immunosuppression in recurrent prostate cancers(7, 8), while combining enzalutamide with checkpoint blockade has not yielded clinical benefits(9). We hypothesize that the association of AR-targeting drugs with recurrent tumor’s immunosuppression are due to selection/adaptation of tumor cells after treatments, rather than a direct immunosuppressive effect. As such, we suggest that to leverage enzalutamide’s potentiating effects on immunotherapies, its direct suppression of PCa cell propagation may be the less important, secondary consideration. Therefore, its regimens (i.e., dosing and treatment time points/durations) may need to be adjusted accordingly. Second, our findings also raise the possibility of repurposing enzalutamide or other AR antagonists for mitigating Treg in immunotherapies, particularly those in which Treg’s immunosuppression is a major barrier to potent immunotherapies, such as glioblastoma(58). Notably, higher AR activity is associated with worse patient survival in glioblastoma(90, 91), and in experimental models, AR signaling promotes GBM cells’ stemness and their resistance to temozolomide, leading to the idea of targeting AR for treating glioblastoma(92–94). However, depletion of androgen (i.e., by castration) has also been found to display intracranial specific, pro-tumorigenic effects in glioblastoma models, suggesting complicated context-dependent effects of AR signaling(95). While the systematic effects of androgen loss remain to be fully elucidated, findings described in this study nevertheless support the idea of targeting AR signaling in T cells for potentiating glioblastoma immunotherapies. We postulate that the strategy may be applicable to both male and female patients for distinct reasons. On one hand, glioblastoma is more prevalent and has a worse prognosis in male patients (who are also more likely to have higher levels of androgens and AR signal activities) (73, 96–100). On the other hand, AR signaling likely exerts similar Treg-suppressing effects in male and female patients, and an AR-targeting therapeutic strategy likely will exert less toxicity in women. Finally, as highlighted by the findings from combining DRP-104 and enzalutamide as described in the current study, it is expected that combinational therapies will be necessary in order to unleash the immunotherapeutic benefits of enzalutamide. While combining with immune checkpoint blockade is an obvious option, other immunotherapeutic approaches such as vaccination-based strategies, exemplified by the anti-Gpc3 vaccination approach described in this study, should also be explored. Further studies testing various combinations and identifying biomarkers for informing novel therapeutic designs are warranted.

## Data availability

The RNA-seq data have been deposited in NCBI’s SRA database and is accessible through SRA accession: PRJNA1447562 and PRJNA1447566.

## Acknowledgements

We thank the Duke Cancer Center Isolation Facility for the support of experiments involving animal models, the Duke Light Microscopy Core Facility for with the support to our imaging experiments, and Dr. Ming Chen (Duke University) for sharing reagents. The quantification of ELISpot results was performed by Duke RBL, which received partial support for construction and renovation from the National Institutes of Health (UC6-AI058607 and G20-AI167200) and facility support from the National Institutes of Health (UC7-AI180254). The work was supported by grants from the National Institutes of Health (5R01-CA260726) and the Prostate Cancer Foundation to J.H. We also acknowledge funding support from The Uncle Kory Foundation to Y.H., The Mike Slive Foundation to C.J.P, and NIH/NCI grant R00 CA237618, the USDA 3092-51000-064-05 and Cancer Prevention and Research Institute of Texas (PR210056, CPRIT) Scholar award in Cancer Prevention and Research to X.G.

## Competing Interests

J.H is a consultant for or owns shares in the following companies: Artera, Kingmed Diagnostics, MoreHealth, OptraScan, York Biotechnology, Chimigen Bio, Pfizer and Sisu Pharma.

## Author Contributions

**Conception and design:** Y He, J Huang

**Methodology development:** J Yu, M Abdulhameed, X Gao, H Staats, CJ Pirozzi, Y He

**Data acquisition:** J Yu, X Jiang, H Yao, C Jin, CJ Pirozzi, X Gao

**Data analysis and interpretation:** J Yu, Z Xing, X Gao, H Staats, Y He, J Huang

**Writing, reviewing, and revision of the manuscript:** J Yu, X Jiang, H Yao, Z Xing, F Zhang, C Jin, H Zhang, M Abdulhameed, ML Bowie, O Meng, DJ George, R Wild, X Gao, Y Zhang, DM Ashley, CJ Pirozzi, H Staats, Y He, J Huang

**Administrative, technical, or material support:** J Yu, H Zhang, X Jiang, F Zhang, M Abdulhameed, ML Bowie, DJ George, R Wild, X Gao, Y Zhang, DM Ashley, CJ Pirozzi, HF Staats, Y He, J Huang

**Study supervision:** Y. He, J Huang

## References

1. Munoz J, Wheler JJ, and Kurzrock R. Androgen receptors beyond prostate cancer: an old marker as a new target. Oncotarget. 2015;6(2):592–603.

2. Guan X, Polesso F, Wang C, Sehrawat A, Hawkins RM, Murray SE, et al. Androgen receptor activity in T cells limits checkpoint blockade efficacy. Nature. 2022;606(7915):791–6.

3. Kwon H, Schafer JM, Song NJ, Kaneko S, Li A, Xiao T, et al. Androgen conspires with the CD8(+) T cell exhaustion program and contributes to sex bias in cancer. Sci Immunol. 2022;7(73):eabq2630.

4. Yang C, Jin J, Yang Y, Sun H, Wu L, Shen M, et al. Androgen receptor-mediated CD8(+) T cell stemness programs drive sex differences in antitumor immunity. Immunity. 2022;55(7):1268–83 e9.

5. Pala L, De Pas T, Catania C, Giaccone G, Mantovani A, Minucci S, et al. Sex and cancer immunotherapy: Current understanding and challenges. Cancer Cell. 2022;40(7):695–700.

6. Lee J, Yurkovetskiy LA, Reiman D, Frommer L, Strong Z, Chang A, et al. Androgens contribute to sex bias of autoimmunity in mice by T cell-intrinsic regulation of Ptpn22 phosphatase expression. Nature communications. 2024;15(1):7688.

7. Consiglio CR, Udartseva O, Ramsey KD, Bush C, and Gollnick SO. Enzalutamide, an Androgen Receptor Antagonist, Enhances Myeloid Cell-Mediated Immune Suppression and Tumor Progression. Cancer Immunol Res. 2020;8(9):1215–27.

8. Xu P, Yang JC, Chen B, Nip C, Van Dyke JE, Zhang X, et al. Androgen receptor blockade resistance with enzalutamide in prostate cancer results in immunosuppressive alterations in the tumor immune microenvironment. J Immunother Cancer. 2023;11(5).

9. Powles T, Yuen KC, Gillessen S, Kadel EE, 3rd, Rathkopf D, Matsubara N, et al. Atezolizumab with enzalutamide versus enzalutamide alone in metastatic castration-resistant prostate cancer: a randomized phase 3 trial. Nature medicine. 2022;28(1):144–53.

10. Xu L, Zhao B, Butler W, Xu H, Song N, Chen X, et al. Targeting glutamine metabolism network for the treatment of therapy-resistant prostate cancer. Oncogene. 2022;41(8):1140–54.

11. Xu L, Yin Y, Li Y, Chen X, Chang Y, Zhang H, et al. A glutaminase isoform switch drives therapeutic resistance and disease progression of prostate cancer. Proceedings of the National Academy of Sciences of the United States of America. 2021;118(13).

12. Moon D, Hauck JS, Jiang X, Quang H, Xu L, Zhang F, et al. Targeting glutamine dependence with DRP-104 inhibits proliferation and tumor growth of castration-resistant prostate cancer. Prostate. 2024;84(4):349–57.

13. Yu J, Jin C, Su C, Moon D, Sun MA, Zhang H, et al. Resilience and Vulnerabilities of Tumor Cells under Purine Shortage Stress. Clinical cancer research : an official journal of the American Association for Cancer Research. 2025;31(20):4345–60.

14. Wang ME, Chen J, Lu Y, Bawcom AR, Wu J, Ou J, et al. RB1-deficient prostate tumor growth and metastasis are vulnerable to ferroptosis induction via the E2F/ACSL4 axis. J Clin Invest. 2023;133(10).

15. Hauck JS, Moon D, Jiang X, Wang ME, Zhao Y, Xu L, et al. Heat shock factor 1 directly regulates transsulfuration pathway to promote prostate cancer proliferation and survival. Commun Biol. 2024;7(1):9.

16. Subramanian A, Tamayo P, Mootha VK, Mukherjee S, Ebert BL, Gillette MA, et al. Gene set enrichment analysis: a knowledge-based approach for interpreting genome-wide expression profiles. Proc Natl Acad Sci U S A. 2005;102(43):15545–50.

17. Mootha VK, Lindgren CM, Eriksson KF, Subramanian A, Sihag S, Lehar J, et al. PGC-1alpha-responsive genes involved in oxidative phosphorylation are coordinately downregulated in human diabetes. Nat Genet. 2003;34(3):267–73.

18. Ge SX, Jung D, and Yao R. ShinyGO: a graphical gene-set enrichment tool for animals and plants. Bioinformatics. 2020;36(8):2628–9.

19. de Leeuw CA, Mooij JM, Heskes T, and Posthuma D. MAGMA: generalized gene-set analysis of GWAS data. PLoS Comput Biol. 2015;11(4):e1004219.

20. Kaya S, Schurman CA, Dole NS, Evans DS, and Alliston T. Prioritization of Genes Relevant to Bone Fragility Through the Unbiased Integration of Aging Mouse Bone Transcriptomics and Human GWAS Analyses. J Bone Miner Res. 2022;37(4):804–17.

21. Morris JA, Kemp JP, Youlten SE, Laurent L, Logan JG, Chai RC, et al. An atlas of genetic influences on osteoporosis in humans and mice. Nat Genet. 2019;51(2):258–66.

22. Tachmazidou I, Hatzikotoulas K, Southam L, Esparza-Gordillo J, Haberland V, Zheng J, et al. Identification of new therapeutic targets for osteoarthritis through genome-wide analyses of UK Biobank data. Nat Genet. 2019;51(2):230–6.

23. Merotto L, Sturm G, Dietrich A, List M, and Finotello F. Making mouse transcriptomics deconvolution accessible with immunedeconv. Bioinform Adv. 2024;4(1):vbae032.

24. He J, Luo H, Wang W, Bu D, Zou Z, Wang H, et al. CIEC: Cross-tissue Immune Cell Type Enrichment and Expression Map Visualization for Cancer. Genomics Proteomics Bioinformatics. 2025;23(1).

25. Vita R, Blazeska N, Marrama D, Members ICT, Duesing S, Bennett J, et al. The Immune Epitope Database (IEDB): 2024 update. Nucleic Acids Res. 2025;53(D1):D436–D43.

26. Schmiedel BJ, Singh D, Madrigal A, Valdovino-Gonzalez AG, White BM, Zapardiel-Gonzalo J, et al. Impact of Genetic Polymorphisms on Human Immune Cell Gene Expression. Cell. 2018;175(6):1701–15 e16.

27. Mathewson ND, Ashenberg O, Tirosh I, Gritsch S, Perez EM, Marx S, et al. Inhibitory CD161 receptor identified in glioma-infiltrating T cells by single-cell analysis. Cell. 2021;184(5):1281–98 e26.

28. Cillo AR, Kurten CHL, Tabib T, Qi Z, Onkar S, Wang T, et al. Immune Landscape of Viral- and Carcinogen-Driven Head and Neck Cancer. Immunity. 2020;52(1):183–99 e9.

29. Chen YP, Yin JH, Li WF, Li HJ, Chen DP, Zhang CJ, et al. Single-cell transcriptomics reveals regulators underlying immune cell diversity and immune subtypes associated with prognosis in nasopharyngeal carcinoma. Cell Res. 2020;30(11):1024–42.

30. Liberzon A, Birger C, Thorvaldsdottir H, Ghandi M, Mesirov JP, and Tamayo P. The Molecular Signatures Database (MSigDB) hallmark gene set collection. Cell Syst. 2015;1(6):417–25.

31. Chalepaki AM, Gkoris M, Chondrou I, Kourti M, Georgakopoulos-Soares I, and Zaravinos A. A multi-omics analysis of effector and resting treg cells in pan-cancer. Comput Biol Med. 2025;189:110021.

32. Kang YJ, Pan L, Liu Y, Rong Z, Liu J, and Liu F. GEPIA3: Enhanced drug sensitivity and interaction network analysis for cancer research. Nucleic Acids Res. 2025;53(W1):W283–W90.

33. Hung MH, Lee JS, Ma C, Diggs LP, Heinrich S, Chang CW, et al. Tumor methionine metabolism drives T-cell exhaustion in hepatocellular carcinoma. Nat Commun. 2021;12(1):1455.

34. Wolf FA, Angerer P, and Theis FJ. SCANPY: large-scale single-cell gene expression data analysis. Genome Biol. 2018;19(1):15.

35. Virtanen P, Gommers R, Oliphant TE, Haberland M, Reddy T, Cournapeau D, et al. SciPy 1.0: fundamental algorithms for scientific computing in Python. Nat Methods. 2020;17(3):261–72.

36. Encarnacion-Rosado J, Sohn ASW, Biancur DE, Lin EY, Osorio-Vasquez V, Rodrick T, et al. Targeting pancreatic cancer metabolic dependencies through glutamine antagonism. Nat Cancer. 2024;5(1):85–99.

37. Pillai R, LeBoeuf SE, Hao Y, New C, Blum JLE, Rashidfarrokhi A, et al. Glutamine antagonist DRP-104 suppresses tumor growth and enhances response to checkpoint blockade in KEAP1 mutant lung cancer. Sci Adv. 2024;10(13):eadm9859.

38. Rais R, Lemberg KM, Tenora L, Arwood ML, Pal A, Alt J, et al. Discovery of DRP-104, a tumor-targeted metabolic inhibitor prodrug. Sci Adv. 2022;8(46):eabq5925.

39. Yokoyama Y, Estok TM, and Wild R. Sirpiglenastat (DRP-104) Induces Antitumor Efficacy through Direct, Broad Antagonism of Glutamine Metabolism and Stimulation of the Innate and Adaptive Immune Systems. Mol Cancer Ther. 2022;21(10):1561–72.

40. Swami U, McFarland TR, Nussenzveig R, and Agarwal N. Advanced Prostate Cancer: Treatment Advances and Future Directions. Trends Cancer. 2020;6(8):702–15.

41. Smith MR, Saad F, Chowdhury S, Oudard S, Hadaschik BA, Graff JN, et al. Apalutamide Treatment and Metastasis-free Survival in Prostate Cancer. N Engl J Med. 2018;378(15):1408–18.

42. Hussain M, Fizazi K, Saad F, Rathenborg P, Shore N, Ferreira U, et al. Enzalutamide in Men with Nonmetastatic, Castration-Resistant Prostate Cancer. N Engl J Med. 2018;378(26):2465–74.

43. de Bono JS, Logothetis CJ, Molina A, Fizazi K, North S, Chu L, et al. Abiraterone and increased survival in metastatic prostate cancer. N Engl J Med. 2011;364(21):1995–2005.

44. Beer TM, Armstrong AJ, Rathkopf DE, Loriot Y, Sternberg CN, Higano CS, et al. Enzalutamide in metastatic prostate cancer before chemotherapy. N Engl J Med. 2014;371(5):424–33.

45. Cao B, Kim M, Reizine NM, and Moreira DM. Adverse Events and Androgen Receptor Signaling Inhibitors in the Treatment of Prostate Cancer: A Systematic Review and Multivariate Network Meta-analysis. Eur Urol Oncol. 2023;6(3):237–50.

46. Cerasuolo M, Maccarinelli F, Coltrini D, Mahmoud AM, Marolda V, Ghedini GC, et al. Modeling Acquired Resistance to the Second-Generation Androgen Receptor Antagonist Enzalutamide in the TRAMP Model of Prostate Cancer. Cancer Res. 2020;80(7):1564–77.

47. Williams CS, Xin Li, Hongjun Jang, Jay Ramanlal Anand, Won Young Lim, Hyejin Lee, Julie Parks, Xingyuan Zhang, Jialiu Xie, Jinshi Zhao, Di Wu, Andrew J. Armstrong, Jessica L. Bowser, Lee Zou, Jiyong Hong, Jason A. Somarelli, Cyrus Vaziri, Pei Zhou. Inhibition of Androgen Receptor Exposes Replication Stress Vulnerability in Prostate Cancer. BioRxiv. 2024.

48. Friedman R. Drug resistance in cancer: molecular evolution and compensatory proliferation. Oncotarget. 2016;7(11):11746–55.

49. Gungabeesoon J, Gort-Freitas NA, Kiss M, Bolli E, Messemaker M, Siwicki M, et al. A neutrophil response linked to tumor control in immunotherapy. Cell. 2023;186(7):1448–64 e20.

50. Hirschhorn D, Budhu S, Kraehenbuehl L, Gigoux M, Schroder D, Chow A, et al. T cell immunotherapies engage neutrophils to eliminate tumor antigen escape variants. Cell. 2023;186(7):1432–47 e17.

51. Faget J, Peters S, Quantin X, Meylan E, and Bonnefoy N. Neutrophils in the era of immune checkpoint blockade. J Immunother Cancer. 2021;9(7).

52. Eruslanov E, Nefedova Y, and Gabrilovich DI. The heterogeneity of neutrophils in cancer and its implication for therapeutic targeting. Nat Immunol. 2025;26(1):17–28.

53. Peyraud F, Guegan JP, Bodet D, Cousin S, Bessede A, and Italiano A. Targeting Tryptophan Catabolism in Cancer Immunotherapy Era: Challenges and Perspectives. Frontiers in immunology. 2022;13:807271.

54. Nishikawa H, and Koyama S. Mechanisms of regulatory T cell infiltration in tumors: implications for innovative immune precision therapies. J Immunother Cancer. 2021;9(7).

55. Curiel TJ. Tregs and rethinking cancer immunotherapy. The Journal of clinical investigation. 2007;117(5):1167–74.

56. Tay C, Tanaka A, and Sakaguchi S. Tumor-infiltrating regulatory T cells as targets of cancer immunotherapy. Cancer cell. 2023;41(3):450–65.

57. Geels SN, Moshensky A, Sousa RS, Murat C, Bustos MA, Walker BL, et al. Interruption of the intratumor CD8(+) T cell:Treg crosstalk improves the efficacy of PD-1 immunotherapy. Cancer cell. 2024;42(6):1051–66 e7.

58. O’Rourke DM, Nasrallah MP, Desai A, Melenhorst JJ, Mansfield K, Morrissette JJD, et al. A single dose of peripherally infused EGFRvIII-directed CAR T cells mediates antigen loss and induces adaptive resistance in patients with recurrent glioblastoma. Science translational medicine. 2017;9(399).

59. Ebert LM, MacRaild SE, Zanker D, Davis ID, Cebon J, and Chen W. A cancer vaccine induces expansion of NY-ESO-1-specific regulatory T cells in patients with advanced melanoma. PloS one. 2012;7(10):e48424.

60. Coretti L, Buommino E, and Lembo F. The aryl hydrocarbon receptor pathway: a linking bridge between the gut microbiome and neurodegenerative diseases. Front Cell Neurosci. 2024;18:1433747.

61. Quintana FJ, and Sherr DH. Aryl hydrocarbon receptor control of adaptive immunity. Pharmacol Rev. 2013;65(4):1148–61.

62. Mezrich JD, Fechner JH, Zhang X, Johnson BP, Burlingham WJ, and Bradfield CA. An interaction between kynurenine and the aryl hydrocarbon receptor can generate regulatory T cells. J Immunol. 2010;185(6):3190–8.

63. Kfoury Y, Baryawno N, Severe N, Mei S, Gustafsson K, Hirz T, et al. Human prostate cancer bone metastases have an actionable immunosuppressive microenvironment. Cancer cell. 2021;39(11):1464–78 e8.

64. Zhao E, Wang L, Dai J, Kryczek I, Wei S, Vatan L, et al. Regulatory T cells in the bone marrow microenvironment in patients with prostate cancer. Oncoimmunology. 2012;1(2):152–61.

65. Sanada N, Gotoh Y, Shimazawa R, Klinge CM, and Kizu R. Repression of activated aryl hydrocarbon receptor-induced transcriptional activation by 5alpha-dihydrotestosterone in human prostate cancer LNCaP and human breast cancer T47D cells. J Pharmacol Sci. 2009;109(3):380–7.

66. Ohtake F, Baba A, Takada I, Okada M, Iwasaki K, Miki H, et al. Dioxin receptor is a ligand-dependent E3 ubiquitin ligase. Nature. 2007;446(7135):562–6.

67. Gandhi VD, Cephus JY, Norlander AE, Chowdhury NU, Zhang J, Ceneviva ZJ, et al. Androgen receptor signaling promotes Treg suppressive function during allergic airway inflammation. The Journal of clinical investigation. 2022;132(4).

68. Altuwaijri S, and Albarrak SM. Androgen Receptor (AR) ablation encumbers the expansion and function of T regulatory cells (Treg) in a male mouse model. Adv Life Sci-Pak. 2025;12(1):197–204.

69. Fijak M, Schneider E, Klug J, Bhushan S, Hackstein H, Schuler G, et al. Testosterone replacement effectively inhibits the development of experimental autoimmune orchitis in rats: evidence for a direct role of testosterone on regulatory T cell expansion. Journal of immunology. 2011;186(9):5162–72.

70. Walecki M, Eisel F, Klug J, Baal N, Paradowska-Dogan A, Wahle E, et al. Androgen receptor modulates Foxp3 expression in CD4+CD25+Foxp3+ regulatory T-cells. Molecular biology of the cell. 2015;26(15):2845–57.

71. Singh RP, and Bischoff DS. Sex Hormones and Gender Influence the Expression of Markers of Regulatory T Cells in SLE Patients. Frontiers in immunology. 2021;12:619268.

72. Lee J, Nicosia M, Hong ES, Silver DJ, Li C, Bayik D, et al. Sex-Biased T-cell Exhaustion Drives Differential Immune Responses in Glioblastoma. Cancer discovery. 2023;13(9):2090–105.

73. Carrano A, Juarez JJ, Incontri D, Ibarra A, and Guerrero Cazares H. Sex-Specific Differences in Glioblastoma. Cells. 2021;10(7).

74. Hu YM, Zhao F, Graff JN, Chen C, Zhang Y, Stommel JM, et al. Elevated Tumor-Associated Androgen Receptor Activity Correlates with Poor Immune Infiltration and Immunotherapy Response across Cancer Types. Cancer Res Commun. 2026;6(1):17–35.

75. Chesner LN, Polesso F, Graff JN, Hawley JE, Smith AK, Lundberg A, et al. Androgen Receptor Inhibition Increases MHC Class I Expression and Improves Immune Response in Prostate Cancer. Cancer discovery. 2025;15(3):481–94.

76. Butler W, Xu L, Zhou Y, Cheng Q, Hauck JS, He Y, et al. Oncofetal protein glypican-3 is a biomarker and critical regulator of function for neuroendocrine cells in prostate cancer. J Pathol. 2023;260(1):43–55.

77. Meng ES, Lim DWT, Chenard-Poirier M, Samol J, Meng RB, Tong Y, et al. Facilitating Selection with Artificial Intelligence-Based Glypican-3 Expression Quantification in Patients with Solid Tumors. Ai Precis Oncol. 2025;2(3):105–16.

78. Yu K, Lin CCJ, Hatcher A, Lozzi B, Kong K, Huang-Hobbs E, et al. PIK3CA variants selectively initiate brain hyperactivity during gliomagenesis. Nature. 2020;578(7793):166–+.

79. Chan ES, Pawel BR, Corao DA, Venneti S, Russo P, Santi M, et al. Immunohistochemical expression of glypican-3 in pediatric tumors: an analysis of 414 cases. Pediatr Dev Pathol. 2013;16(4):272–7.

80. Sawada Y, Yoshikawa T, Nobuoka D, Shirakawa H, Kuronuma T, Motomura Y, et al. Phase I trial of a glypican-3-derived peptide vaccine for advanced hepatocellular carcinoma: immunologic evidence and potential for improving overall survival. Clin Cancer Res. 2012;18(13):3686–96.

81. Sawada Y, Sakai M, Yoshikawa T, Ofuji K, and Nakatsura T. A glypican-3-derived peptide vaccine against hepatocellular carcinoma. Oncoimmunology. 2012;1(8):1448–50.

82. Brown AT, and Wagner C. Regulation of enzymes involved in the conversion of tryptophan to nicotinamide adenine dinucleotide in a colorless strain of Xanthomonas pruni. J Bacteriol. 1970;101(2):456–63.

83. Tagawa ST. Neuroendocrine prostate cancer after hormonal therapy: knowing is half the battle. Journal of clinical oncology : official journal of the American Society of Clinical Oncology. 2014;32(30):3360–4.

84. Seguier D, Parent P, Duterque-Coquillaud M, Labreuche J, Fromont-Hankard G, Dariane C, et al. Emergence of Neuroendocrine Tumors in Patients Treated with Androgen Receptor Pathway Inhibitors for Metastatic Prostate Cancer: A Systematic Review and Meta-analysis. Eur Urol Oncol. 2025;8(2):581–90.

85. Beltran H, and Demichelis F. Therapy considerations in neuroendocrine prostate cancer: what next? Endocr Relat Cancer. 2021;28(8):T67–T78.

86. Deng X, Elzey BD, Poulson JM, Morrison WB, Ko SC, Hahn NM, et al. Ionizing radiation induces neuroendocrine differentiation of prostate cancer cells in vitro, in vivo and in prostate cancer patients. American journal of cancer research. 2011;1(7):834–44.

87. Li Y, He Y, Butler W, Xu L, Chang Y, Lei K, et al. Targeting cellular heterogeneity with CXCR2 blockade for the treatment of therapy-resistant prostate cancer. Science translational medicine. 2019;11(521).

88. Chen Z, Cai A, Zheng H, Huang H, Sun R, Cui X, et al. Carbidopa suppresses prostate cancer via aryl hydrocarbon receptor-mediated ubiquitination and degradation of androgen receptor. Oncogenesis. 2020;9(5):49.

89. Sun F, Indran IR, Zhang ZW, Tan MH, Li Y, Lim ZL, et al. A novel prostate cancer therapeutic strategy using icaritin-activated arylhydrocarbon-receptor to co-target androgen receptor and its splice variants. Carcinogenesis. 2015;36(7):757–68.

90. Farina-Jeronimo H, Martin-Ramirez R, Gonzalez-Fernandez R, Medina L, de Vera A, Martin-Vasallo P, et al. Androgen deficiency is associated with a better prognosis in glioblastoma. Eur J Med Res. 2024;29(1):57.

91. Farina-Jeronimo H, de Vera A, Medina L, and Plata-Bello J. Androgen Receptor Activity Is Associated with Worse Survival in Glioblastoma. J Integr Neurosci. 2022;21(3):86.

92. Méndez ABD, Di Giuliani M, Sacconi A, Tremante E, Lulli V, Di Martile M, et al. Androgen receptor inhibition sensitizes glioblastoma stem cells to temozolomide by the miR-1/miR-26a-1/miR-487b signature mediated WT1 and FOXA1 silencing. Cell Death Discov. 2025;11(1).

93. Zalcman N, Canello T, Ovadia H, Charbit H, Zelikovitch B, Mordechai A, et al. Androgen receptor: a potential therapeutic target for glioblastoma. Oncotarget. 2018;9(28):19980–93.

94. Zhao X, Li Y, Martin WR, Salem F, Korem R, Bolbol S, et al. Blood-Brain Barrier (BBB)-Penetrable Androgen Receptor (AR) Degrader as a Potential Therapeutic Agent for Glioblastoma. ACS Pharmacol Transl Sci. 2026;9(5):1164–76.

95. Lee J, Chung YM, Silver DJ, Hao Y, Harwood DSL, Ealy A, et al. Androgen loss accelerates brain tumour growth via HPA axis activation. Nature. 2026.

96. Yang W, Warrington NM, Taylor SJ, Whitmire P, Carrasco E, Singleton KW, et al. Sex differences in GBM revealed by analysis of patient imaging, transcriptome, and survival data. Science translational medicine. 2019;11(473).

97. Goldberg M, Frank LS, Altawalbeh G, Negwer C, Wagner A, Gempt J, et al. Do clinical outcomes in individuals with malignant gliomas differ between sexes? Brain Spine. 2025;5.

98. Ostrom QT, Rubin JB, Lathia JD, Berens ME, and Barnholtz-Sloan JS. Females have the survival advantage in glioblastoma. Neuro-oncology. 2018;20(4):576–7.

99. Leppert J, Ditz C, Matschke J, Krenzlin H, Keric N, Ziemann C, et al. Sex-related Survival Differences in Patients With Glioblastoma - Results From a Retrospective Analysis. Anticancer Research. 2025;45(5):2059–69.

100. Jovanovich N, Habib A, Chilukuri A, Hameed NUF, Deng H, Shanahan R, et al. Sex-specific molecular differences in glioblastoma: assessing the clinical significance of genetic variants. Front Oncol. 2023;13:1340386.

