## Supplemental figures for "A Glutamine antagonist-modulated tumor microenvironment unleashes enzalutamide’s immunotherapeutic effects"

### Supplementary Figures

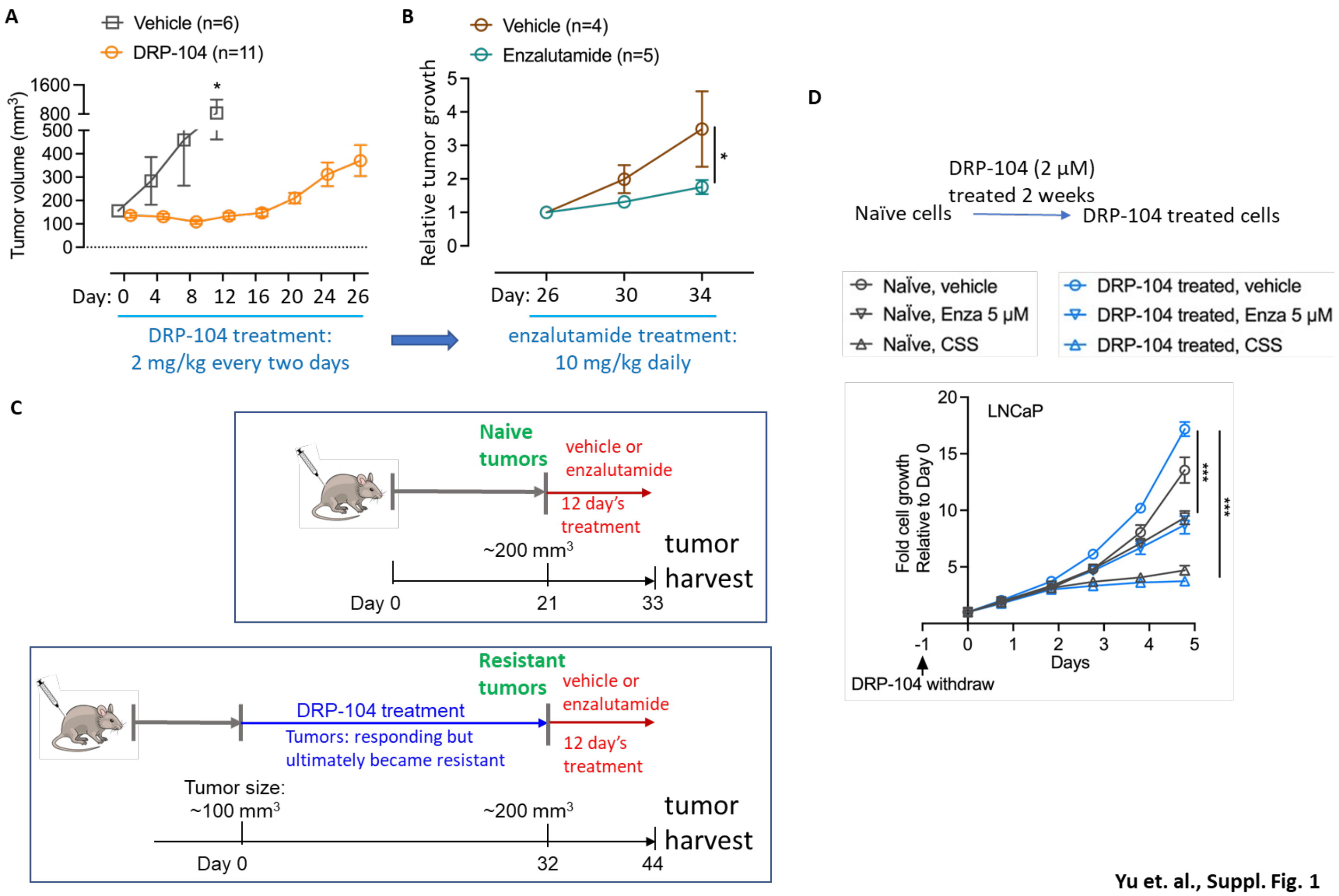


**Yu et. al., Suppl. Fig. 1**

**Suppl. Fig. 1. DRP-104 resistant tumors display apparent vulnerability to enzalutamide in immune competent hosts. (A)** TrampC2 cells were used to generate subcutaneous tumors in wildtype C56BL/6 host mice. Tumors (~100-200 $mm^3$ in size) were treated with DRP-104 following the indicated regimen, and their sizes were measured every two days. **(B)** Tumors that became resistant to DRP-104 (indicated by their apparent progression between day 16 to day 26 at which pointes their sizes were ~300-400 $mm^3$) were subsequently treated with vehicle control or with enzalutamide following the indicated regimen and tumor sizes were determined. **(C)** Generation of TrampC2-derived naïve tumors and DRP-104-resistant tumors and the subsequent treatments (with vehicle control or with enzalutamide). **(D)** Naïve or DRP-104-treated (2 μM for two weeks) LNCaP cells were subjected to treatment with enzalutamide (enza) or charcoal dextran stripped fetal bovine serum (CSS) cultured condition, and relative cell propagation was determined via Incucyte. Data are depicted as mean ± SEM. *p<0.05, ***p<0.001.

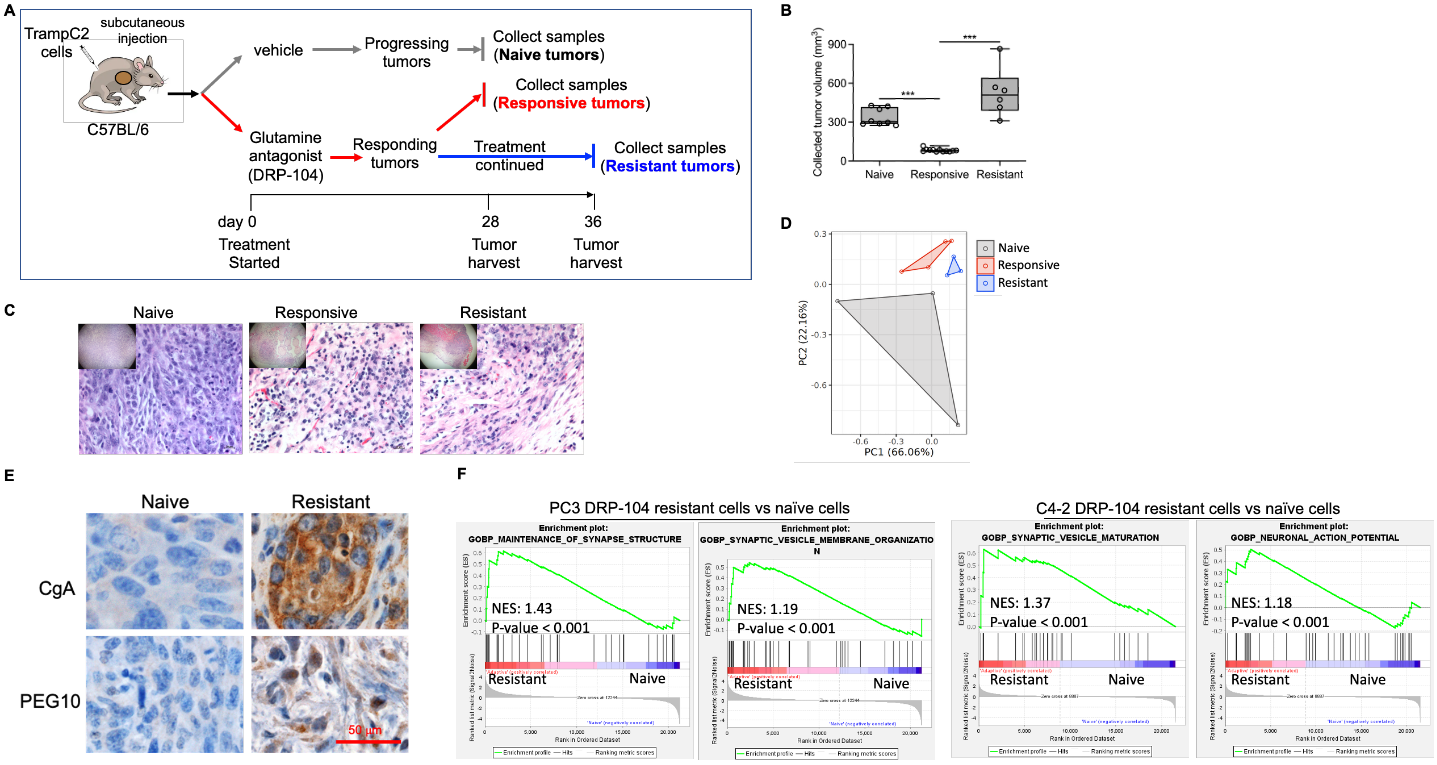


Yu et. al., Suppl. Fig. 2

**Suppl. Fig. 2. Distinct features of DRP-104-native (untreated), responsive and resistant tumors. (A)** Generation of TrampC2-derived naïve tumors, DRP-104-responsive tumors and DRP-104-resistant tumors. **(B)** Tumor volumes at the times of tumor harvest. **(C)** Harvested tumor tissues were subjected to H&E staining and representative images were shown. **(D)** mRNA-seq results were subjected to principal component analysis (PCA). **(E)** Naïve and Resistant tumors were used for staining against CgA and PEG10 and representative images were shown. **(F)** Matched pairs of DRP-104 resistant versus parental human PCa cell lines (PC3 and C4-2) were subjected to mRNA-seq and detected genes in the resistant cells (versus the parental (naive) cells were used for pathway enrichment analysis. ***p<0.001.

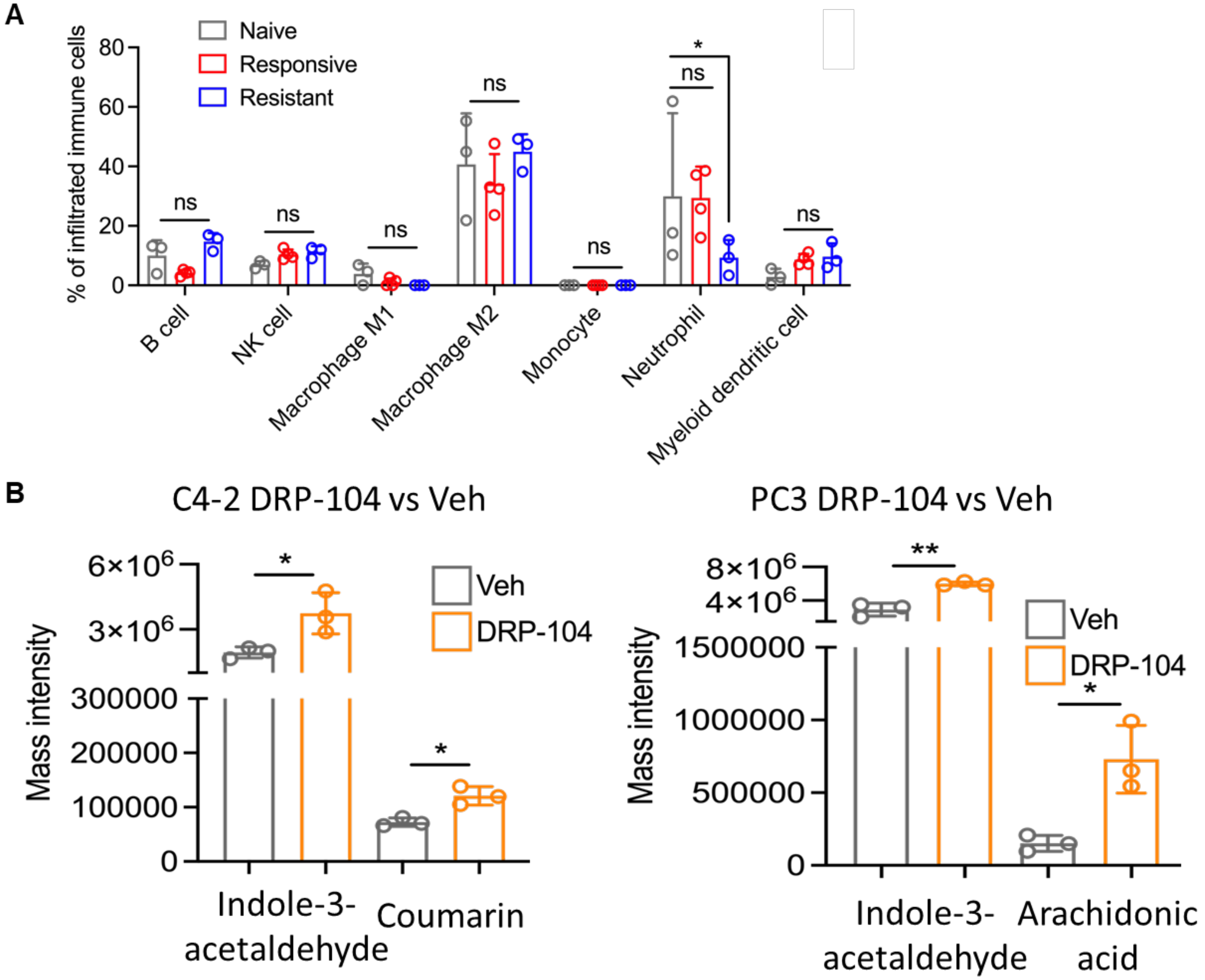


**Yu et. al., Suppl. Fig. 3**

**Suppl. Fig. 3. DRP-104 treatment modulates tumor immune microenvironment and alters the abundance of immunosuppressive metabolites. (A)** mRNA-seq data from tumors described in Fig. 2 (A-B) were used for cell deconvolution analyses to determine the compositions of the cells in the three groups of tumors. The estimated fractions various types of immune cells in these three groups of tumors were shown. **(B)** Relative abundance of indicated metabolites in human PCa cell lines (PC3 and C4-2) that were treated with vehicle control, or with DRP-104 was shown (the metabolite profiling data was obtained from our recently described dataset). *p<0.05, **p<0.01. ns: no significant.

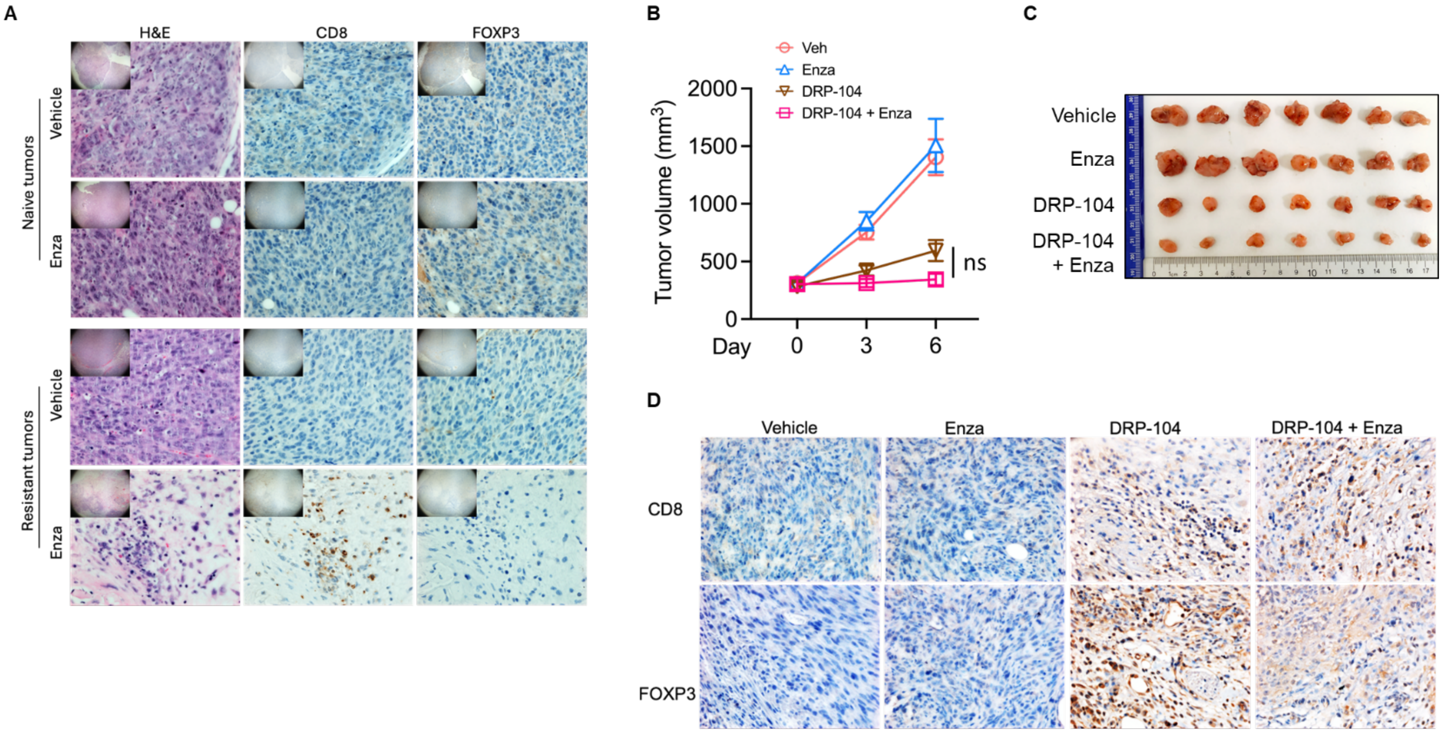


Yu et. al., Suppl. Fig. 4

**Suppl. Fig. 4. Combining DRP-104 and enzalutamide results in superior therapeutic efficacy and depletes Treg from tumors. (A)** DRP-104 naïve or resistant tumors were treated with vehicle control or with enzalutamide for 12 days, at which point most vehicle-treated DRP-104-resistant tumors reached the humane endpoint (tumor volumes of ~1000 $mm^3$), and all tumors were harvested were IHC analysis. The representative images were shown (Enza: enzalutamide). **(B-C)** C57BL/6 mice bearing subcutaneous tumors derived from PPR, a mouse NEPC-like cell line, were used for four arms of treatments: vehicle control, Enza monotherapy (10 mg/kg) every day, DRP-104 (2 mg/kg) every two days, or both agents, and the tumor growth was determined. The growth curves (B) and images of tumors harvested at the human endpoint (C) were shown. **(D)** Tumors shown in Fig. 4G were used for anti-CD8 and anti-FOXP3 IHC staining and the representative IHC images were shown. ns: no significant.

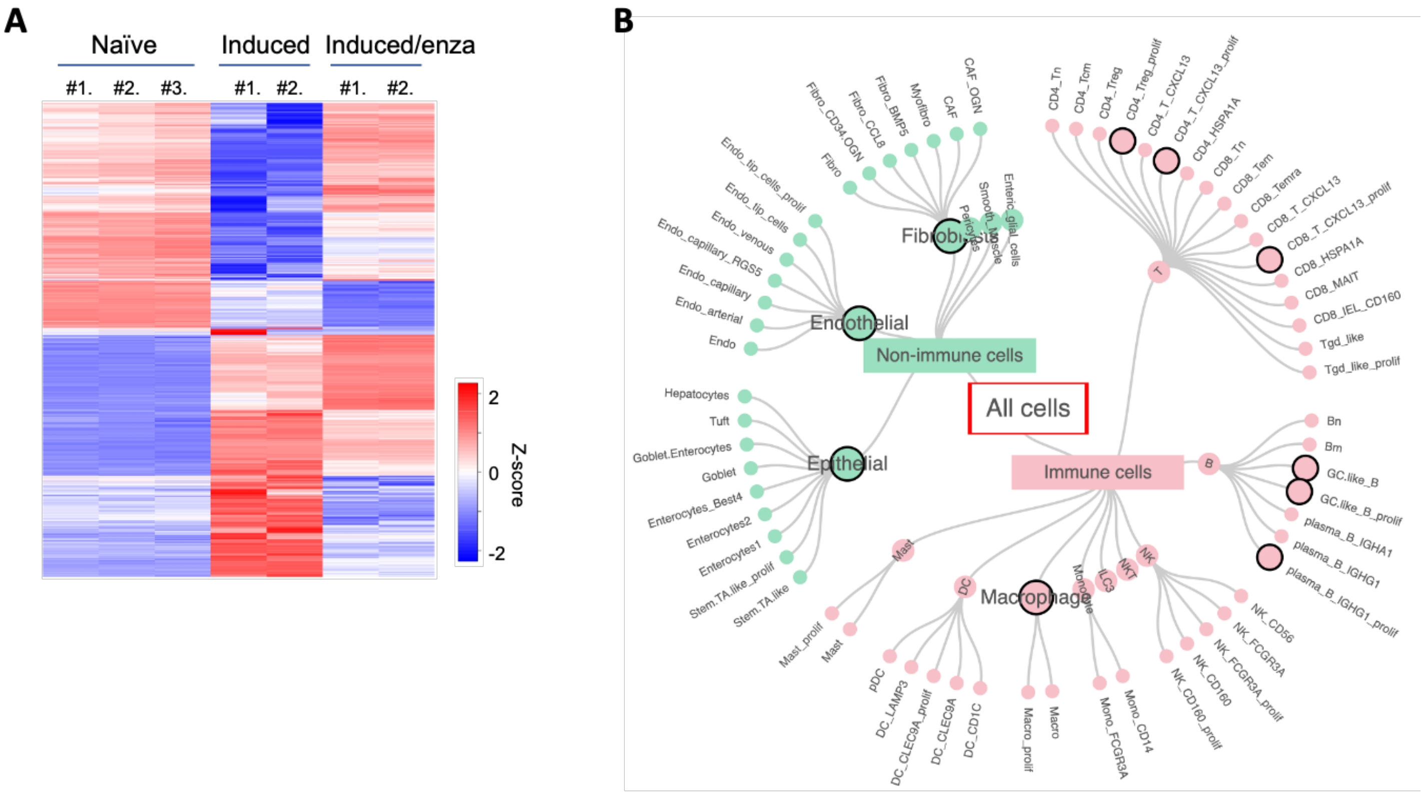


Yu et. al., Suppl. Fig. 5

**Suppl. Fig. 5. Effects of enzalutamide on the transcriptomic profiles of splenocyte-derived, CD4-enriched cells undergoing Treg induction. (A)** A heat map demonstrating the effects of enzalutamide on DEGs identified from comparing Induced versus Naïve cells (Padj<0.05, n=1155 genes, including log2FC>1 / 607 genes and log2FC<-0.5/ 548 genes). **(B)** Genes whose expression was affected by Treg induction (Induced cells versus Naïve cells; Padj<=0.25) and found to be altered by enzalutamide (Induced/enza cells versus Induced cells; Padj<=0.25) were used for CIEC analysis to estimate immune cell types/states (cell types/states marked by black circles indicated those that were significantly enriched).

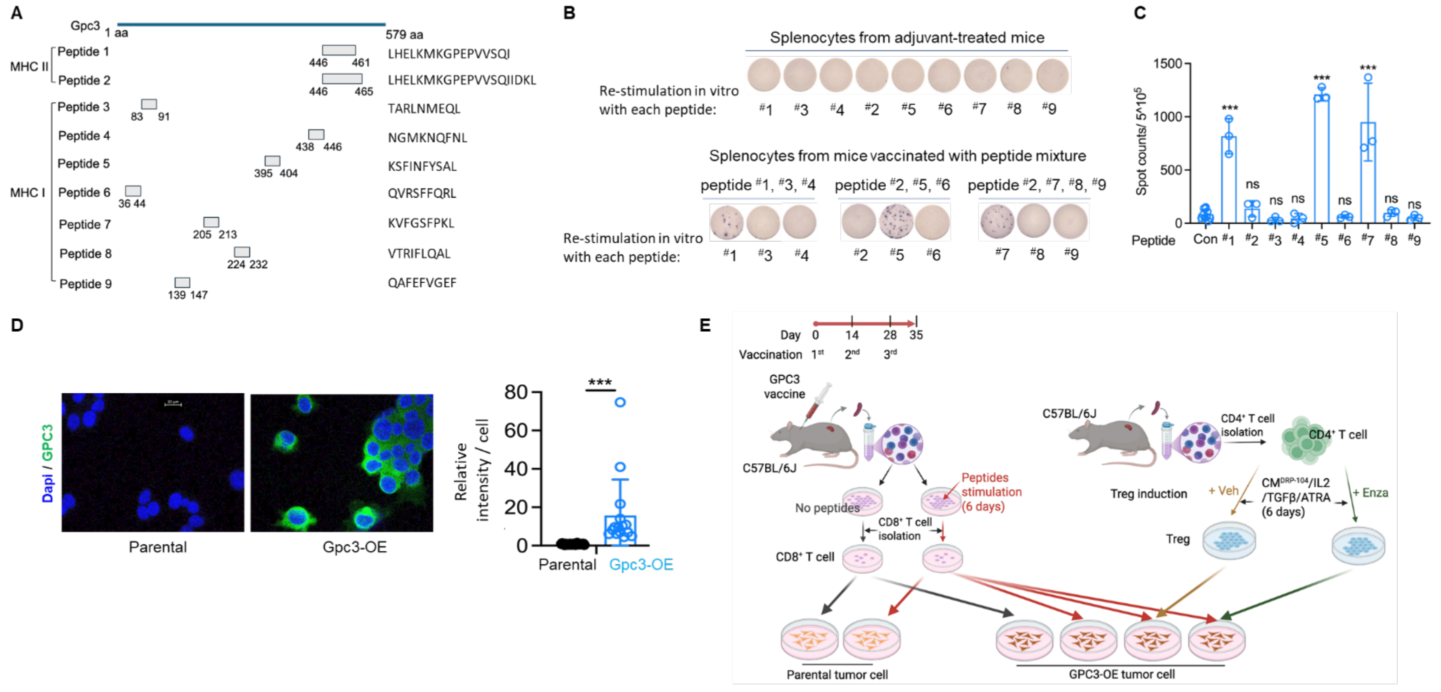


Yu et. al., Suppl. Fig. 6

**Suppl. Fig. 6. An anti-GPC3 peptide vaccination model and its utilization for CTL assays in vitro. (A)** Nine mouse Gpc3-derived peptides predicted to be presented by MHC-I or MHC–II were used for vaccination to test their immunogenicity in vivo. **(B-C)** Splenocytes from mice vaccinated with adjuvants or with each of the three peptide mixtures were used for in vitro re-stimulation each indicated peptide, and anti-IFNɣ ELISPot assay was performed. Representative images (B) and quantifications (C) were shown. **(D)** Parental TrampC2 or its derivative line with exogenous Gpc3 overexpression (Gpc3-OE) were used for anti-Gpc3 IF staining, and the representative images and quantification were shown. **(E)** A scheme shows the mouse vaccination, cytotoxic T cell preparation, the Treg differentiation, and the CTL assays. ***p<0.001.
